# Genome and tissue-specific transcriptome of the tropical milkweed (*Asclepias curassavica*)

**DOI:** 10.1101/2024.01.12.575411

**Authors:** Honglin Feng, Jing Zhang, Adrian F. Powell, Gretta L. Buttelmann, Lily Yang, Ethan Yan, Fumin Wang, Steven B. Broyles, Georg Jander, Susan R. Strickler

## Abstract

Tropical milkweed (*Asclepias curassavica*) serves as a host plant for monarch butterflies (*Danaus plexippus*) and other insect herbivores that can tolerate the abundant cardiac glycosides that are characteristic of this species. Cardiac glycosides, along with additional specialized metabolites, also contribute to the ethnobotanical uses of *A. curassavica*. To facilitate further research on milkweed metabolism, we assembled the 197 Mbp genome of a fifth-generation inbred line of *A. curassavica* into 619 contigs, with an N50 of 10 Mbp. Scaffolding resulted in 98% of the assembly being anchored to 11 chromosomes, which are mostly colinear with the previously assembled common milkweed (*A. syriaca*) genome. Assembly completeness evaluations showed that 98% of the BUSCO gene set is present in the *A. curassavica* genome assembly. The transcriptomes of six tissue types (young leaves, mature leaves, stems, flowers, buds, and roots), with and without defense elicitation by methyl jasmonate treatment, showed both tissue-specific gene expression and induced expression of genes that may be involved in cardiac glycoside biosynthesis. Expression of a *CYP87A* gene, the predicted first gene in the cardiac glycoside biosynthesis pathway, was observed only in the stems and roots and was induced by methyl jasmonate. Together, this genome sequence and transcriptome analysis provide important resources for further investigation of the ecological and medicinal uses of *A. curassavica*.

## Introduction

Plants in the genus *Asclepias* (milkweeds) are known for their production of cardiac glycosides, which function as inhibitors of essential Na^+^/K^+^-ATPases in animal cells (Kreis and Müller-Uri, 2010; Agrawal, 2017). These defensive compounds are found the roots, leaves, flowers, seeds, latex and other parts of *Asclepias* plants (López-Goldar et al., 2022; Betz et al., 2024). Specialized herbivores from at least six insect orders have developed target-site resistance to the cardiac glycosides in their *Asclepias* hosts (Dobler et al., 2012; Petschenka et al., 2017; Karageorgi et al., 2019). In particular, the iconic interactions between milkweeds and *Danaus plexippus* (monarch butterflies), which sequester toxic cardiac glycosides from their milkweed host plants for their own protection, have been the subject of extensive ecological research and hundreds of publications.

Among the more than 200 known milkweed species, *Asclepias syriaca* (common milkweed) and *Asclepias curassavica* (tropical milkweed; Figure 1A) are the most common host plants for monarch caterpillars (Agrawal, 2017). Whereas *A. syriaca* is found in temperate areas of North America, the native range of *A. curassavica* comprises tropical North and South America, including the Caribbean islands (Woodson, 1954). A shrubby perennial, *A. curassavica* has been introduced as a garden ornamental or a naturalized weed in other tropical areas of the world. In areas where *A. syriaca* does not grow, *A. curassavica* is often the main host plant for monarch butterflies. For instance, caterpillars from non-migratory monarch populations in southern Florida were reported to feed primarily on introduced *A. curassavica* (Knight and Brower, 2009). The monarch butterfly Na^+^/K^+^-ATPase, which is resistant to most cardiac glycosides, is inhibited by voruscharin (Figure 1B), a sulfur and nitrogen-containing cardiac glycoside that constitutes about 40% of the total cardiac glycosides in *A. curassavica* (Agrawal et al., 2021).

**Figure 1.**
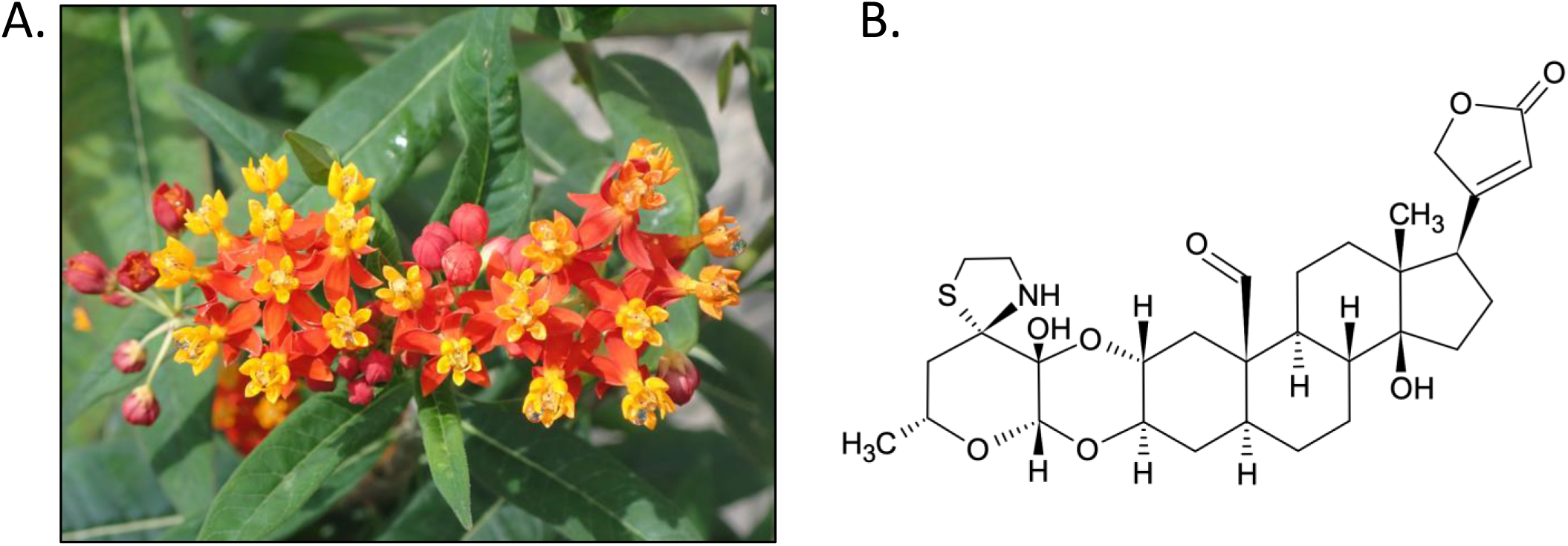
*Asclepias curassavica*, tropical milkweed. A. Flowers of the greenhouse-grown *A. curassavica l*ineage that was used for the current study. B. Structure of voruscharin, the most abundant cardiac glycoside in *A. curassavica*.

Due to its ethnobotanical uses, Carl Linnaeus named the genus *Asclepias* after Asclepius, the Greek god of medicine (Linnaeus, 1754; Quattrocchi, 1999). The numerous reported medical applications of *A. curassavica* include the antimicrobial (Navarro Garcia et al., 2003; Peraza-Sanchez et al., 2007; Jacobo-Salcedo Mdel et al., 2011; Alonso-Castro et al., 2021) and anticancer (Yuan et al., 2016; Zheng et al., 2019) properties of plant extracts. More broadly, cardiac glycosides produced by milkweed and other plants, in particular foxglove (*Digitalis* spp.), have been used for centuries to treat congestive heart failure (Tomov, 1981; Rehman and Hai, 2023). The first enzyme of the cardiac glycoside biosynthesis pathway, a cytochrome P450 that catalyzes sterol side chain cleavage, has been confirmed in species from three plant families: Plantaginaceae, woolly foxglove (*Digitalis lanata*) and common foxglove (*Digitalis purpurea*); Apocynaceae, Sodom apple (*Calotropis procera*); and Brassicaceae, wormseed wallflower (*Erysimum cheiranthoides*) (Carroll et al., 2023; Kunert et al., 2023; Younkin et al., 2024).

Although *A. syriaca* genome assemblies have been described previously (Straub et al., 2011; Weitemier et al., 2019; Boyle et al., 2022), this species is not very tractable as a laboratory model system. In contrast to *A. syriaca*, *A. curassavica* grows well in pots in the greenhouse, does not require a winter diapause, can be propagated from cuttings, and is self-fertile (Wyatt and Broyles, 1997). To facilitate molecular genetic research on the extensive ecological interactions and medical applications of *Asclepias*, we initiated an *A. curassavica* genome sequencing project. Here we present the assembly and annotation of a high-quality genome, as well as the transcriptomes of six tissue types, from a fifth-generation inbred lineage of *A. curassavica*.

## Methods

### Plants and growth conditions

*Asclepias curassavica* seeds were initially purchased from Butterfly Encounters (www.butterflyencounters.com/). Plants were grown in a greenhouse under ambient light and were self-pollinated manually for five generations. To produce a larger number of seeds in the sixth generation, a flowering plant was placed in a cage with a bumblebee colony (*Bombus terrestris*; https://www.biobestgroup.com) for more efficient pollination. Seeds from the sixth generation of inbreeding were submitted to the USDA-ARS Ornamental Plant Germplasm Center (https://opgc.osu.edu/ ; https://www.ars-grin.gov/ ; accession number OPGC 7586). The fifth-generation inbred line of *A. curassavica* was used to prepare material for genomic DNA and cDNA sequencing.

For plant growth, seeds were washed three times with 0.6% sodium hypochlorite in deionized water with one drop of dish soap. After the washes, seeds were transferred to a 90 mm Petri dish containing a layer of moist brown paper towel. Sealed Petri dishes with seeds were incubated at 30°C for 3-6 days, until germination. After the seeds had germinated, seedlings were moved to pots containing Cornell Mix (by weight 56% peat moss, 35% vermiculite, 4% lime, 4% Osmocote slow-release fertilizer [Scotts, Marysville, OH], and 1% Unimix [Scotts]) in Conviron (Winnipeg, Canada) growth chambers with a 16:8 photoperiod, 180 µM m^-2^ s^-1^ photosynthetic photon flux density, 60% humidity, and constant 23 °C temperature.

### Methyl jasmonate treatment

Synchronized flowering *A. curassavica* plants were treated with methyl jasmonic acid before tissue collection. Each plant was sprayed with 0.01% methyl jasmonate (dissolved in 0.1% ethanol) to cover the whole plant, both sides of the leaves and the stem, to the point of liquid run-off, in a separated spray room. Control plants were sprayed with 0.1% ethanol. After the spray treatment, methyl jasmonate-treated and control-treated plants were separated in two different greenhouses with the same temperature and light settings. After 24 hours, all plants were sprayed again. Six hours after the booster spray, different plant tissues, including young leaves (the top 4 newly emerged leaves), mature leaves, stems, flowers, buds, and roots, were collected, flash-frozen in liquid nitrogen, and stored at -80°C until use. Unopened flowers (buds) and open flowers (Figure1) were collected as separate tissue types, but no effort was made to define specific bud developmental stages or time after flower opening for tissue collection. The six tissue types were collected in eight-fold replication, with and without methyl jasmonate treatment, for a total of 96 samples.

### DNA extraction, library preparation, and sequencing

To extract intact DNA for long-read sequencing, *A. curassavica* leaf tissue was harvested and placed into 2 ml screw-cap vials, 3 three-mm steel balls were added to each tube, tubes were flash-frozen in liquid nitrogen, and samples were homogenized on a paint shaker (Harbil, Wheeling, IL, USA). DNA was extracted from the homogenized tissue using a modified cetyl trimethylammonium bromide (CTAB) protocol using chloroform:isoamyl alcohol 24:1 (Fulton et al., 1995). To ensure that DNA fragments were of the correct size for Nanopore sequencing, the DNA was then size-selected using the Circulomics SRE kit v2.0 (SS-100-101-01, PacBio, Menlo Park, CA, USA). A concentration of 150 ng/µL of DNA was initially used, and 55 µL of Buffer EB was added prior to the incubation step. To prepare the DNA sequencing library, the Oxford Nanopore Genomic DNA by Ligation kit (SQK-LSK109, Oxford Nanopore Technologies, New York, NY, USA) was used with the undiluted ∼2.5 mg of SRE-treated DNA from the previous step. Sequencing was done using a Nanopore MinION flow cell (flow cell number FAL75881-1282) on a MinION sequencing device. To maximize the available runtime of the flow cell and generate more data, the library was split in two and each half was sequenced successively on the same flow cell, one before and one after completing the Nanopore Nuclease flush protocol (Oxford Nanopore Technologies, New York, NY, USA).

For Illumina sequencing, DNA was extracted from *A. curassavica* leaves using the Wizard® Genomic DNA Purification Kit (Promega, Madison WI, USA). The quantity and quality of genomic DNA was assessed using a Qubit 3 fluorometer (Thermo Fisher, Waltham, MA, USA) and a Bioanalyzer DNA12000 kit (Agilent, Santa Clara, CA, USA). The PCR-free TruSeq DNA (Illumina, San Diego, CA) was used to prepare a library. Samples were sequenced on an Illumina MiSeq instrument (paired-end 2×150 bp) at the Cornell University Biotechnology Resource Center (Ithaca, NY, USA). A Hi-C library was prepared from *A. curassavica* leaves using the Proximo Hi-C Plant Kit (Phase Genomics, Seattle, WA, USA) and was sequenced on an Illumina MiSeq instrument (paired-end 2×75bp) at the Cornell University Biotechnology Resource Center (Ithaca, NY, USA).

### RNA extraction, cDNA library preparation, and sequencing

RNA samples were prepared using the RNeasy® Plant Mini Kit (Qiagen, Hilden, Germany) according to the manufacturer’s protocol. Briefly, tissue samples were ground using a 1600 MiniG® Automated Tissue Homogenizer and Cell Lyser (Cole-Parmer, Vernon, IL, USA) and ∼100 mg of each sample were used for RNA extraction. RNA concentrations were measured using a Qubit® RNA BR kit (ThermoFisher Scientific, Waltham, MA, USA) on a Qubit® 3.0 Fluorometer (ThermoFisher Scientific), and 10 samples were randomly selected and evaluated using the Agilent 2100 bioanalyzer (Agilent, Santa Clara, CA, USA) to ensure high RNA quality (RNA Integrity Number > 7).

High-quality RNA samples were used for cDNA library preparation with the Illumina Stranded mRNA Prep kit following a ligation protocol (Illumina, San Diego, CA, USA). Briefly, from the total RNAs (∼ 1 ug), we first captured and enriched mRNA with polyA tails using oligo (dT) magnetic beads. Purified mRNA samples were then fragmented and denatured for first-strand cDNA synthesis using the first-strand synthesis master mix with reverse transcriptase.

Then, the mRNA templates were removed, and second-strand cDNA was synthesized to generate blunt-ended double-stranded cDNA fragments. In place of deoxythymidine triphosphate (dTTP), deoxyuridine triphosphate, (dUTP) was incorporated to quench the second strand during amplification to achieve strand specificity. The cDNA fragments were captured and purified using the Agencourt AMPure XP magnetic beads (Beckman Coulter, Brea, CA, USA). The blunt-ended double-stranded cDNA was adenylated at the 3’ end to prevent self-ligation and facilitate ligation to the adapters with corresponding thymine (T) nucleotides on the 3’ end in a later step. Adenylated cDNA fragments were further ligated with pre-index RNA anchors for dual indexing. To amplify cDNA libraries, the anchor-ligated DNA fragments were selectively amplified and dual-indexed using unique indexes and primer sequences with 10 cycles of PCR. Finally, the cDNA libraries were cleaned up using AMPure XP magnetic beads and concentrations were measured using the Qubit® dsDNA BR Assay kit (ThermoFisher Scientific) using a Qubit® 3.0 Fluorometer.

RNAseq data were generated using Illumina sequencing at the Cornell University Biotechnology Resource Center (Ithaca, NY). The 96 uniquely dual-indexed cDNA libraries were pooled based on their concentrations measured using a Qubit® 3.0 Fluorometer. The pooled libraries were first evaluated using the Agilent 2100 Bioanalyzer for quality check and then sequenced with the Illumina MiSeq platform (Illumina) to ensure approximately equal amounts input of each individual library for Illumina HiSeq. Finally, about 5 million 2×75 bp pair-ended RNAseq reads were generated for each of the 96 libraries using the Illumina NextSeq500 platform (Illumina).

### Genome assembly and annotation

Genome size was estimated by k-mer analysis (K=19, K = 21, and K=29) of Illumina reads using Jellyfish v2.2.7 (Marçais and Kingsford, 2011) and GenomeScope.v1.0 (Vurture et al., 2017). Genomic *A. curassavica* nanopore and Illumina sequences were assembled using MaSurCA v3.4.1 (Zimin et al., 2013). The assembly was corrected using Illumina sequences and Pilon v1.23 (Walker et al., 2014). Redundancy was removed using Purge Haplotigs (Roach et al., 2018). Hi-C was used to scaffold the contigs using 3D-DNA v180922 (Dudchenko et al., 2017). The assembly was manually adjusted in Juicebox v2.15.07 (Durand et al., 2016). Assembly gaps were filled using the nanopore reads and draft assembly as input to LR_gapcloser v3 (Xu et al., 2019). quarTeT version 1.2.1 (Lin et al., 2023) was used for telomere and centromere analysis.

LTR_retriever (Ou and Jiang, 2018) was used with outputs from LTRharvest (Ellinghaus et al., 2008) and LTR_FINDER (Xu and Wang, 2007) to identify long terminal repeat retrotransposons (LTRs). The LTR library was then used to hard mask the genome, and RepeatModeler version: open-1.0.11 (Smit and Hubley, 2015) was used to identify additional repetitive elements in the remaining unmasked segments of the genome. Protein-coding sequences were excluded using blastx v2.8.1+ (Altschul et al., 1990; Ellinghaus et al., 2008) results in conjunction with the ProtExcluder.pl script from the ProtExcluder v1.2 package (Campbell et al., 2014). The libraries from RepeatModeler and LTR_retriever were then combined and used with RepeatMasker v4.0.9 (Smit et al., 2015) to produce the final masked version of the genome.

RNAseq reads were mapped to the genome with HISAT2 v2.1.0 (Kim et al., 2015). Portcullis v1.2.0 (Mapleson et al., 2018) and Mikado v2.3.2 (Venturini et al., 2018) were used to process and filter the resulting bam files. Augustus v3.3.3 (Stanke et al., 2008) and Snap v2006-07-28 (Korf, 2004) were trained and implemented through the Maker v2.31.10 pipeline (Cantarel et al., 2008), with proteins from Swiss-Prot (Boutet et al., 2007) and processed RNAseq added as evidence. Gene models were filtered with the following criteria: 1) at least one match found in the Trembl database (4-17-19) (Boutet et al., 2007) with an E-value less than 1e-20, 2) InterProScan matches to repeats were removed, 3) genes with an AED score of 1 and no InterPro domain were removed, and 4) single-exon genes with no InterPro domain were removed. Functional annotation and classification were performed using BLASTx v2.8.1+ (Altschul et al., 1990) and InterProScan v5.36-75.0 (Jones et al., 2014). Both genome and annotation completeness were assessed by BUSCO v3.1.0 (Waterhouse et al., 2017) using the embryophyta lineage set.

### Synteny, gene family, and phylogenetic analysis

A synteny plot between the *A. curassavica* and *A. syriaca* pseudomolecules was created using sequence matches calculated with nucmer v4.0.0beta2 (Kurtz et al., 2004) and plotted using Circos v0.69-6 (Krzywinski et al., 2009). OrthoFinder v2.5.2 (Emms & Kelly, 2015) was used to identify gene families using protein sequences from genome-sequenced close relatives to *A. curassavica* with genome annotations. The following species were included: *A. curassavica* v1.0*, Asclepias syriaca* v1.0 (Boyle et al., 2022), *Calotropis gigantea* (Hoopes et al., 2018), *Catharanthus roseus* (Kellner et al., 2015), *Rhazya stricta* (Sabir et al., 2016), *Eustoma grandiflorum* (Liang et al., 2022), *Gelsemium sempervirens* (Frank et al., 2015), *Neolamarckia cadamba* (Zhao et al., 2022), *Chiococca alba* (Lau et al., 2020), *Coffea canephora* v1.0 (Denoeud et al., 2014). These results were used to create an Upset plot using the UpSetR package v1.4.0 (Conway et al., 2017) depicting shared and unique gene clusters. Maximum likelihood methods were used to identify significantly expanded and contracted gene families in *Asclepias*. Gene families from Orthofinder and the species tree were input into CAFE v.3.0 (Han et al., 2013). Before inputting, the tree was ultrametricized with penalized likelihood using the chronopl() function in the R package APE (Paradis et al., 2004). An error model and a single lambda rate, a measure of the rate of evolutionary change in the tree calculated by the CAFE v. 3.0 software (Han et al., 2013), were determined for the tree. The species phylogeny built by Orthofinder was dated using treePL (Farrell et al., 2001) by calibrating to the MRCAs of the UC Conservative Ages crown_treePL estimates of the Apocynaceae and Rubiaceae estimated in (Ramírez-Barahona et al., 2020). For the gene families showing significant expansion and contraction (p<0.05) in *A curassavica*, we retrieved the list of associated gene ontology (GO) terms from our annotation. The resulting list of expanded or contracted *A. curassavica* orthogroups and their associated GO terms was input to topGO (Alexa and Rahnenfuhrer, 2023) for enrichment analyses. We searched for enrichment in GO terms associated with biological functions, molecular process, and cellular component and used Fisher’s exact test to determine significance.

For phylogenetic analysis of the CPY87A4 gene family, the *D. lanata* CPY87A4 protein sequence (Accession ID OR134564) was retrieved from Genbank. The gene family that best matched this protein based on sequence identity was identified by BLAST queries and the alignment was retrieved from the OrthoFinder multiple sequence alignment results. Additional sequences were added to the alignment, focusing on five cardiac glycoside-producing species, *A. curassavica, A. syriaca, C. procera, C. gigantea,* and *E. cheiranthoides*, as well as the well-studied model plant species *Nicotiana benthamiana, Solanum lycopersicum, Arabidopsis thaliana,* and *Oryza sativa.* The protein sequences in FASTA format are in Supplemental Dataset S3. Clustal Omega (Sievers et al., 2011) was used to generate the sequence alignment in Figure S5. A maximum likelihood, midpoint-rooted tree was created using IQ-Tree and 1,000 replicates for calculating bootstrap values (Hoang et al., 2018; Minh et al., 2020). A machine-readable (Newick Format) version of the tree is in Supplementary Dataset S4. The tree in Figure 9 was visualized using MEGA11 (Tamura et al., 2021).

### Differential expression analysis

The paired-end sequencing reads from cDNA libraries were aligned to gene models from the reference *A. curassavica* genome using bowtie v2.2.5 with default settings (Langmead and Salzberg, 2012). The raw read counts for each gene model were extracted using the pileup.sh script from BBMap v35.85 (https://sourceforge.net/projects/bbmap/). Raw read counts were then used to generate a data matrix for differential expression analyses using the Bioconductor package edgeR v3.36.0 (Robinson et al., 2010) in Rstudio v2022.02.3 (https://www.rstudio.com). For differential expression analyses, the raw read counts for each gene were first normalized as counts per million (CPM) to the total read counts of all genes in each sample. Only genes with expression levels > 2 CPM and detected in at least 8 samples were retained for further analyses. The filtered normalized reads were then investigated using principal component analysis (PCA) to exclude the possibility of outliers or batch effects. Then, differential expression analyses were performed between the methyl jasmonate and control treatments, independently within each tissue type. To identify differentially expressed genes (methyl jasmonate vs control), we applied the exactTest function followed by the decideTestsDGE function in edgeR, where we retained genes with a significant differential expression value of p ≤ 0.05, and a Benjamini & Hochberg adjusted false discovery rate (FDR) ≤ 0.05 (Robinson et al., 2010).

To facilitate visualization of variations in gene expression, the common differentially expressed genes across different tissue types were identified using a web-based Venn diagram generator from Bioinformatics & Evolutionary Genomics (https://bioinformatics.psb.ugent.be) and the result was plotted with UpSetR v1.4.0 (Conway et al., 2017). To understand the function of those genes, in addition to using the existing genome annotation, we further annotated the genes by blasting their protein sequences against Uniprot database, using PANNZER2 (Törönen et al., 2018) and local BLAST (Altschul et al., 1997; Schäffer et al., 2001), as well as KEGG pathway annotation (https://www.genome.jp/kegg/pathway.html). To visualize the expression results, we generated a heatmap for all the log2 fold changes of the common differentially expressed genes using the R package pheatmap v1.0.12 (Kolde, 2018).

## Results

### A. curassavica genome assembly and annotation

Based on flow cytometry, the *A. curassavica* genome was estimated to have a 639 Mbp diploid size (Lewis and Ruter, 2023). We used paired-end Illumina and Oxford Nanopore sequencing, with 93X and 45X coverage, respectively, to assemble a diploid genome from a fifth-generation inbred line of *A. curassavica*. K-mer analysis using values of K ranging from 19 to 29 suggested that the haploid genome size is 161 Mbp, which is smaller than the previous estimate, with a heterozygosity of 0.2% (Figure S1). The Illumina and nanopore sequencing reads were assembled into 851 contigs, with an N50 of 2.4 Mbp and a total length of 241 Mbp (Table S1). Duplicate removal, scaffolding, and gap filling produced a final assembly with 633 contigs, with an N50 of 10 Mbp and a 197 Mbp total assembly length (Table S1). Approximately 98% of the final assembly was anchored to 11 chromosomes (Figure S2). Telomeres and centromeres were not reliably identified in the assembly. Assembly completeness evaluations using BUSCO showed that 98% of the BUSCO gene set is present in the assembly, and only 0.9% of the genes are duplicated (Table S1).

Repeat analysis indicates that 51.5% of the genome is composed of repetitive elements, with LTR elements being the most abundant (Figure 2). Gene prediction resulted in 26,035 gene models, which tended to be located toward distal regions of the chromosomes (Figure S3). The annotation captures 95.2% of the BUSCO set (Table S1). Altogether, 21,221 genes were functionally annotated and 13,196 were assigned GO terms.

**Figure 2:**
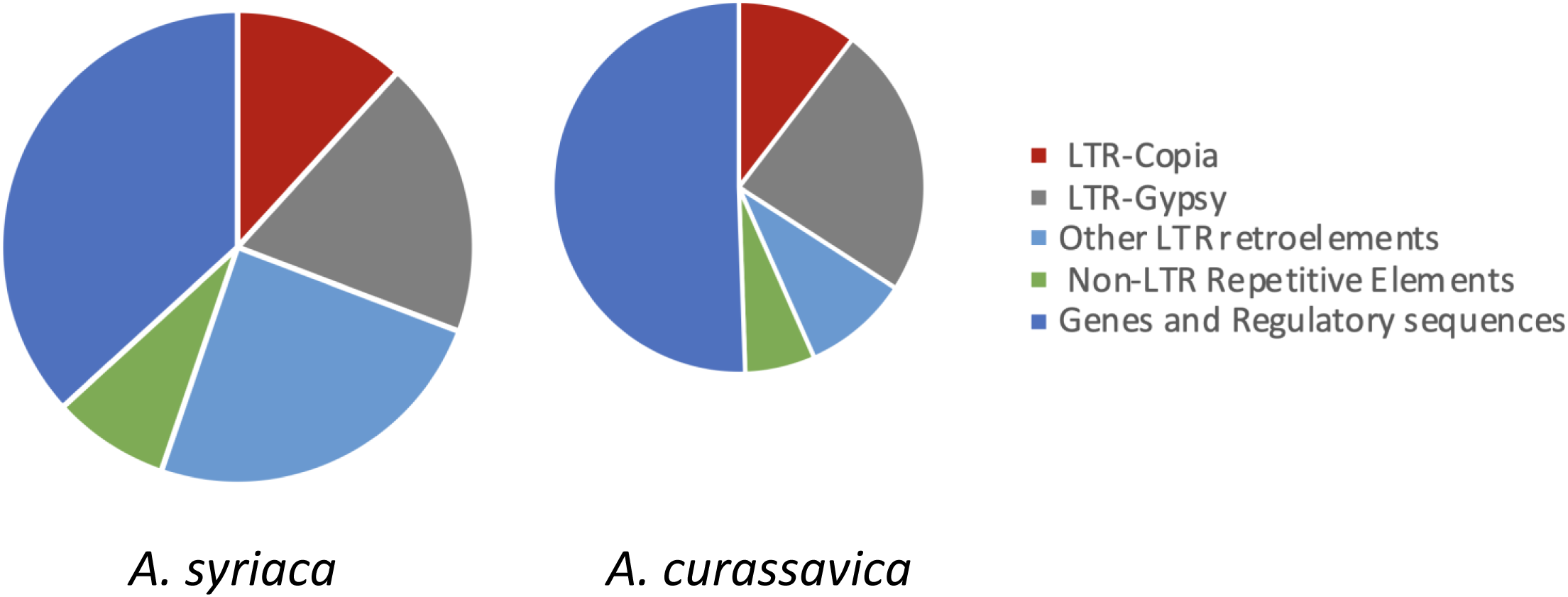
Pie charts showing percentages of genome features in the *Asclepias syriaca and Asclepias curassavica* pseudomolecules. The size of each pie is scaled to the genome size of the species.

### Evolution of the A. curassavica genome

The 11 pseudomolecules of the *A. curassavica* genome assembly are highly colinear with the *A. syriaca* genome assembly (Boyle et al., 2022) (Figure 3), although there are a few fusions and breakpoints, *e.g*., on chromosomes 1, 10, and 11. The *A. syriaca* genome was found to have a greater expansion of LTR retroelements relative to *A. curassavica*, contributing to its larger size (Figure 2). Contracted areas of the *A. curassavica* genome relative to *A. syriaca* are generally in regions that contain noncoding and repetitive regions. Gene family analysis involving closely related species with sequenced genomes, *A. curassavica, A. syriaca*, *C. gigantea*, *C. roseus*, *C. alba*, *C. canephora, E. grandiflorum*, *G. sempervirens*, *M. speciosa, N. cadamba*, and *R. stricta*, assigned 92.5% of detected genes (336,215/ 363,631) to 26,866 orthogroups. A total of 9,052 gene families containing 168,416 genes were shared between all species (Figure 4). To identify gene families that had expanded or contracted we first estimated the lambda rate, a measure of the rate of evolutionary change, as 0.638 using the CAFE software package. A total of 406 gene families have expanded in *A. curassavica* and 564 have contracted (Figure 5). We then focused on rapidly evolving gene families that had changed significantly along the *A. curassavica* lineage. Rapidly expanding gene families in *A. curassavica* include genes involved in amine metabolic processes and defense responses, whereas rapidly contracting families included genes involved in magnesium ion transport and manganese ion homeostasis (Table S2). We did not observe a significant expansion or contraction of the *CYP87A* gene family, which has been associated with cardiac glycoside biosynthesis in other plant species (Carroll et al., 2023; Kunert et al., 2023; Younkin et al., 2024).

**Figure 3:**
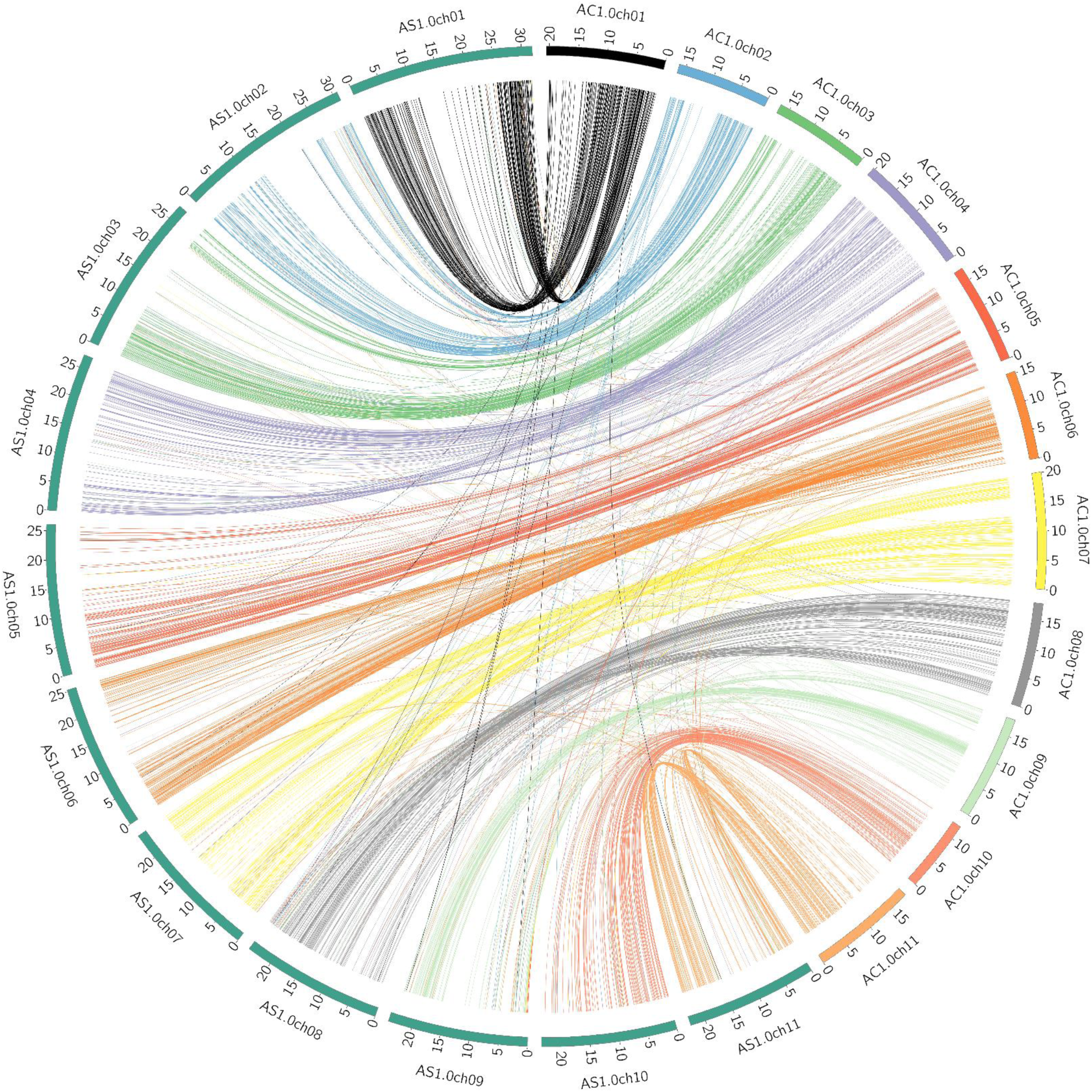
Synteny plot showing regions of collinearity between the *Asclepias curassavica* and *Asclepias syriaca* pseudomolecules. Sequence matches were calculated with nucmer and plotted using Circos. Numbers next to the chromosome arcs indicate Mbp lengths.

**Figure 4:**
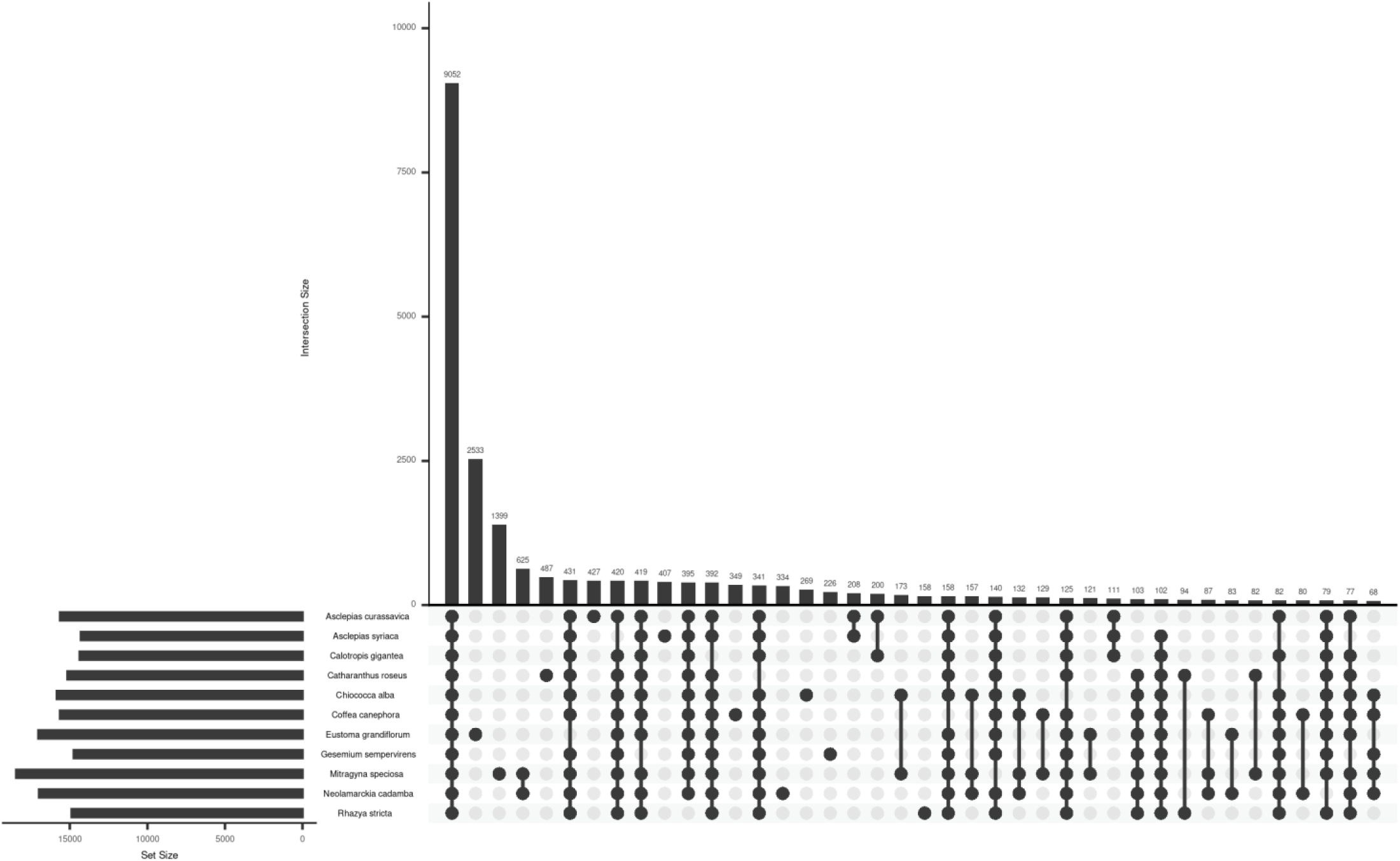
Upset plot showing the intersection of gene families among *Asclepias curassavica* and sequenced near relatives. The numbers of gene families are indicated for each species and species intersection.

**Figure 5:**
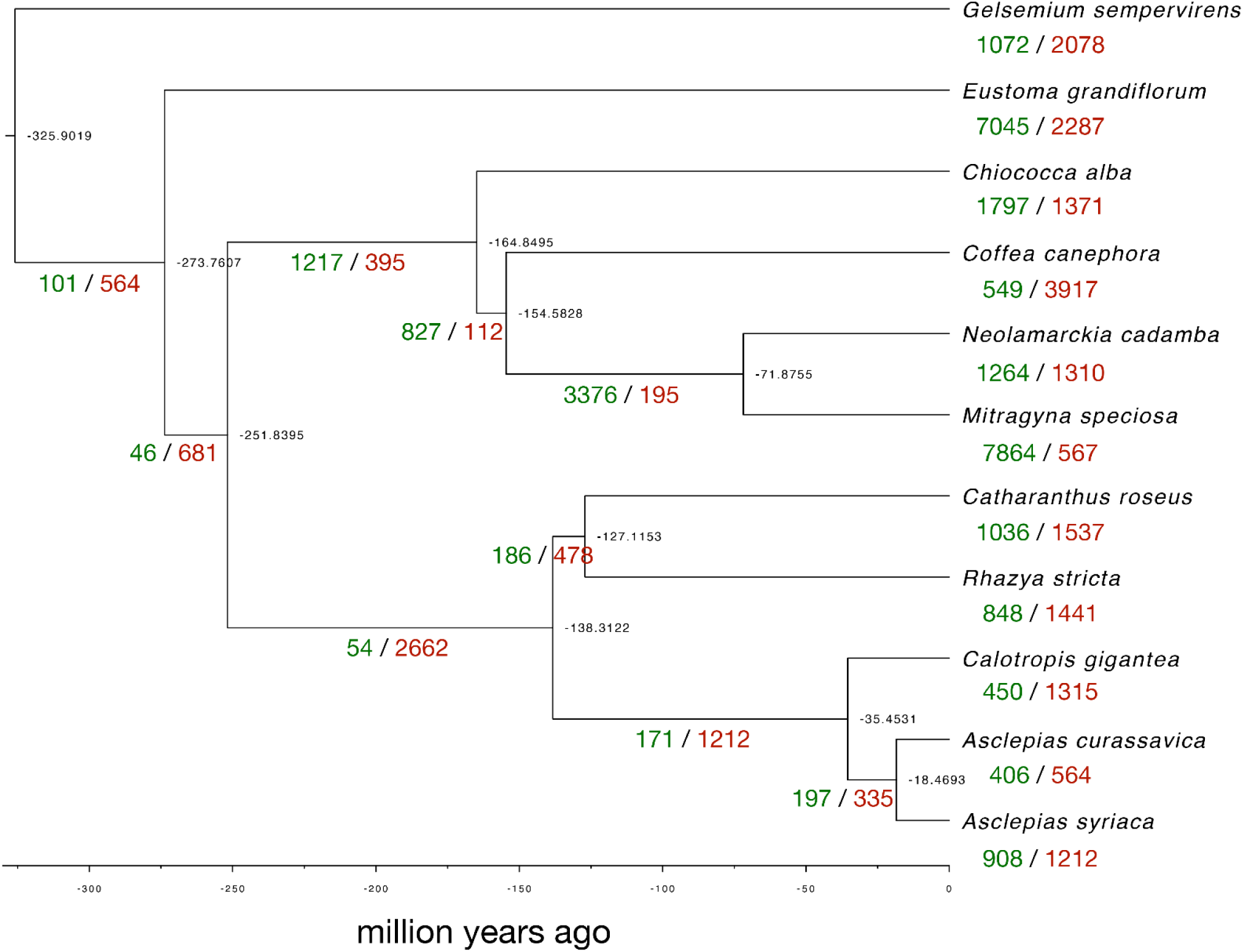
Species phylogeny showing gene family expansions and contractions. Expansions are show in green and contractions are shown in red. Estimated divergence times (millions of years before the present) are in black at the nodes of the tree.

### Differentially expressed genes upon jasmonic acid treatment

The expression of plant genes varies among tissue types and in response to environmental variation. For instance, insect herbivory elicits the production of defenses and other metabolic changes in plants. This induction, which depends on jasmonate-dependent signaling pathways, can be elicited by treatment with methyl jasmonate (Howe and Jander, 2008). To increase the diversity of the expressed genes that can be used for genome annotation, we collected samples from six *A. curassavica* tissue types (buds, roots, stems, flowers, young leaves, and mature leaves), with and without methyl jasmonate treatment, in 8-fold replication.

Illumina sequencing (RNAseq) generated approximately 5 million paired-end reads for each of the *A. curassavica* samples. Approximately 75% of RNAseq reads of each sample were mapped to the coding sequences of gene models annotated in the genome. Relative gene expression levels are presented in Supplemental Datasets S1 and S2. Principal component analysis (PCA) showed that the overall gene expression patterns from both control samples (Figure 6A) and methyl jasmonate -treated samples (Figure 6B) were distinct in each tissue type. Similarly, PCA clustering based on tissue types demonstrated clear separation between control and methyl jasmonate-treated samples (Figure 6C-H). It is worth noting that, compared to other tissue types, samples from buds showed less distinct separation (Figure 6C), possibly due to their rapid growth and dynamic changes in gene expression in this tissue type.

**Figure 6.**
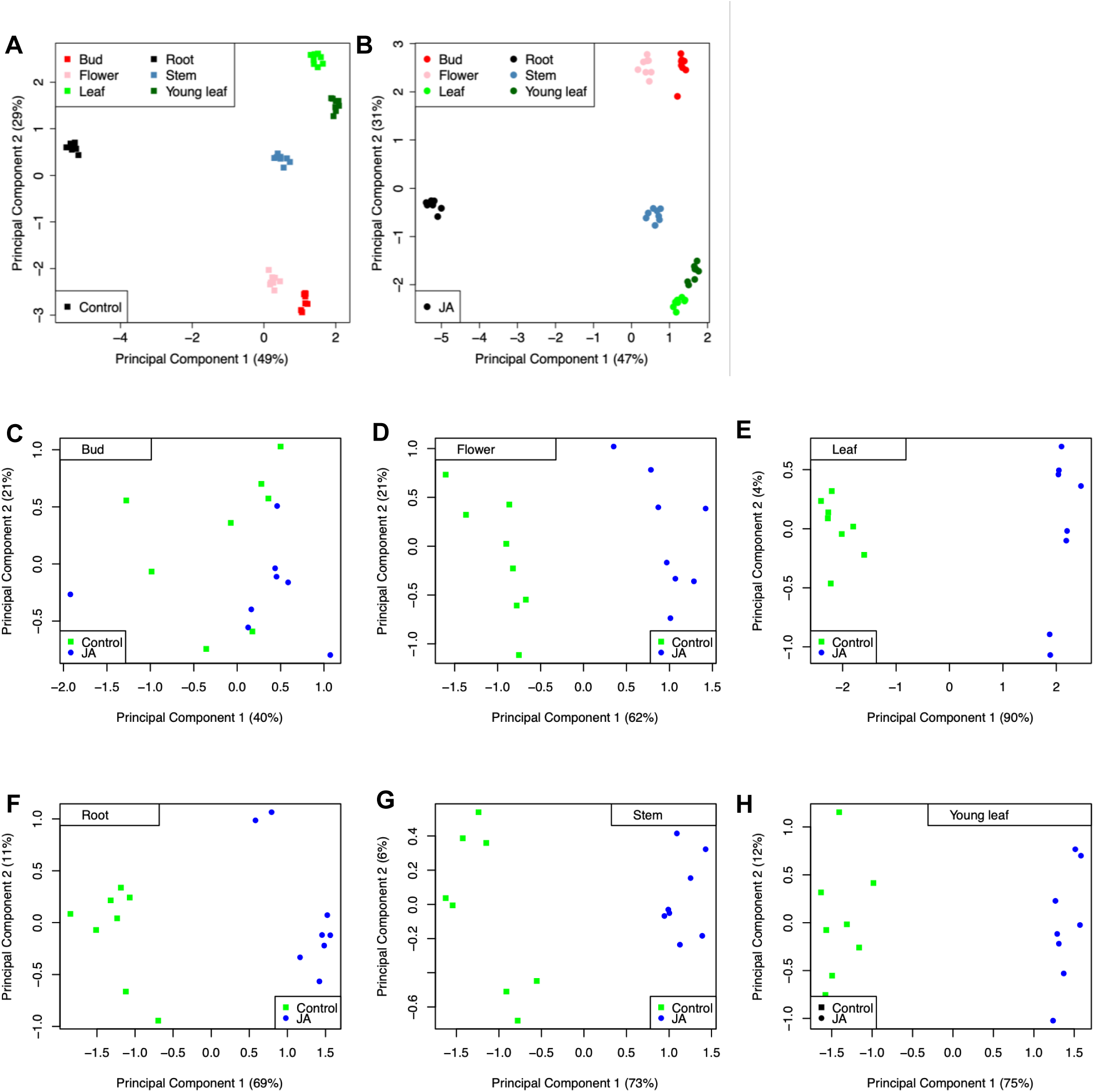
Principal component analyses (PCA) of RNAseq data. (A) PCA of control samples treated with 0.1% ethanol (B) PCA of methyl jasmonate-treated samples. (C-H) comparison of control samples (green dots) and methyl jasmonate-treated samples (blue dots) of (C) buds, (D) flowers, (E) mature leaves, (F) roots, (G) stems, and (H) young leaves. Gene expression data are in Supplemental Datasets S1 and S2.

Differential gene expression analyses were conducted independently for each tissue type comparing methyl jasmonate-treated samples to controls. In buds, roots, stems, flowers, young leaves, and mature leaves, we identified 2465, 4329, 5864, 6262, 7769, and 9608 differentially expressed genes, respectively (Figure 7). While many of these differentially expressed genes were specific to individual tissue types (blue bars in Figure 7), 349 were differentially expressed in a similar manner across all tissue types (red bar in Figure 7). Functional annotation resulted in 89% (*i.e*., 310/349) of the differentially expressed genes being assigned putative functions (Figure S4 and Table S3).

**Figure 7.**
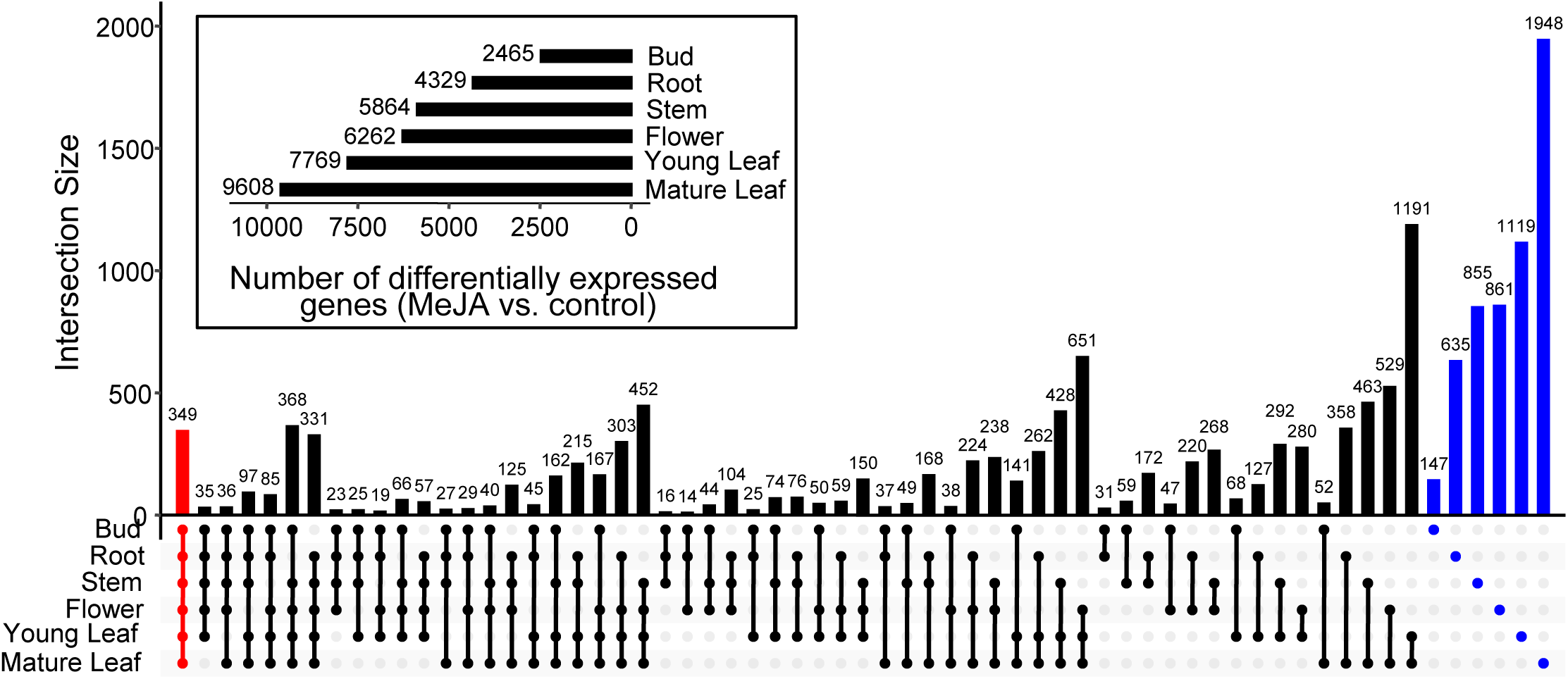
Number of differentially expressed genes across tissue types after methyl jasmonic acid (MeJA) treatment. The horizontal bar plot in the inset box shows the number of differentially expressed genes (MeJA-treated *vs*. control) in each of the six tested tissue types. The intersection plot shows the number of differentially expressed genes across different tissue type combinations. The red bar represents the number of core differentially expressed genes across all tissue types, blue bars represent the number of differentially expressed genes that are specific to each tissue type, and black bars represent other tissue intersections indicated by the dot connections under the bar plot. This plot was generated using UpSetR v1.4.0. Gene expression data are in Supplemental Datasets S1 and S2.

Specific gene expression changes indicated that methyl jasmonate treatment elicited a robust defense response in *A. curassavica.* Across all six tissue types, 18 genes associated with jasmonic acid biosynthesis and signaling were consistently upregulated upon methyl jasmonate treatment (Figure 8 and Table S3) (Ruan et al., 2019; Li et al., 2022; Hewedy et al., 2023). These genes included five *JAZ* (jasmonate-ZIM domain) genes, which play a crucial role in the jasmonate signaling cascade, as well as *ACOX1* (acyl-coezyme A oxidase 1), *AOC* (allene oxide cyclase), *AOS* (allene oxide synthase), two *LOX* (linoleate 13S-lipoxygenases) genes, *KAT2* (3-ketoacyl-CoA thiolase 2), *4CLL5* (4-coumarate-CoA ligase-like 5), *NINJA* (novel interactor of JAZ), and *MYC2* (a jasmonate-responsive transcription factor). In addition, methyl jasmonate treatment induced other genes associated with plant defense responses, including *ITH* (inhibitor of trypsin and hageman factor), a proteinase inhibitor that is potentially involved in plant resistance to herbivory (Chen et al., 1992), as well as *PAMP-A70* (pathogen-associated molecular patterns-induced protein A70) and *R1A-10* (late blight resistance protein homolog), which are both associated with plant disease resistance (Figure 8 and Table S3).

**Figure 8.**
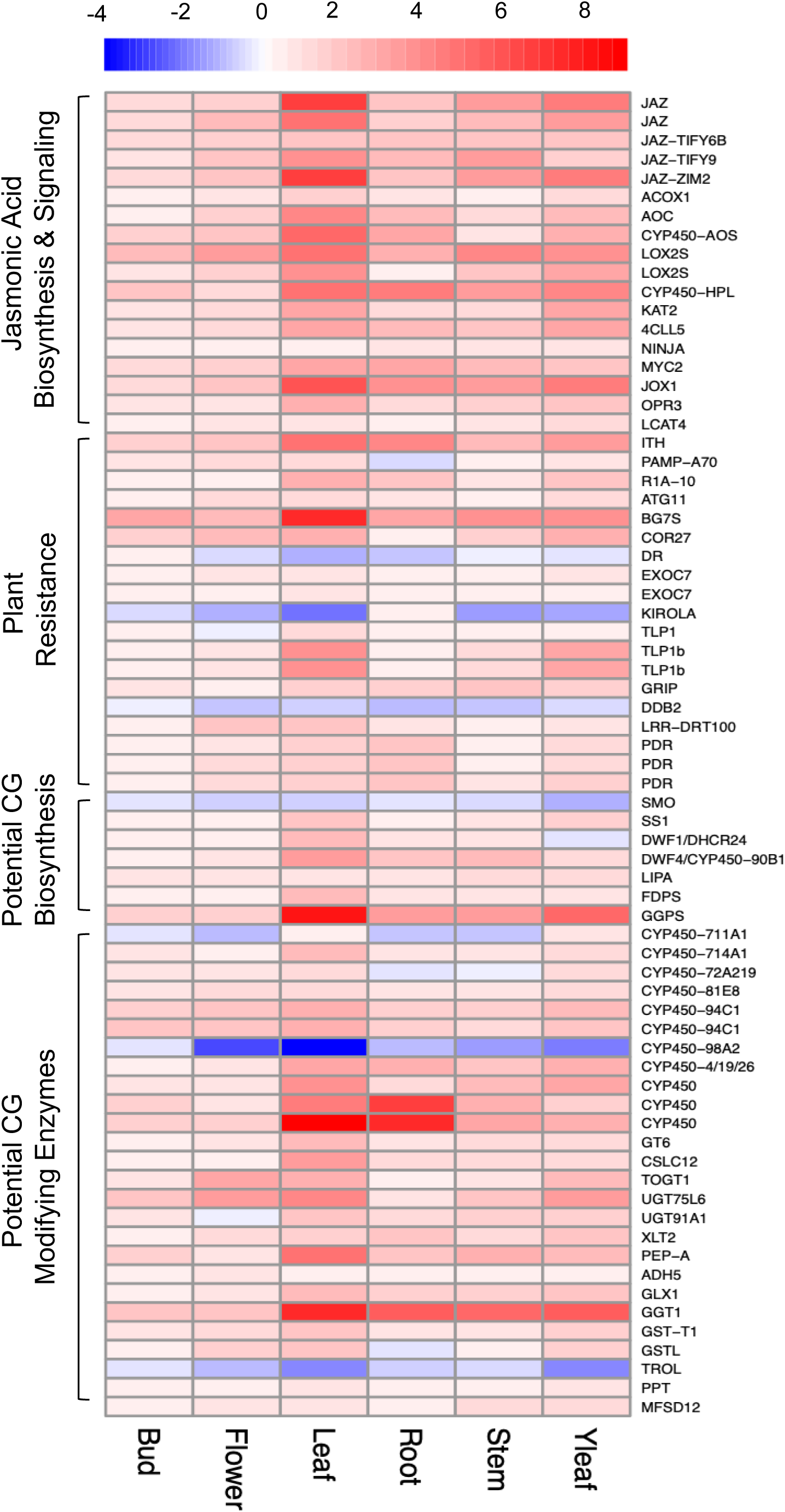
Heatmap of log_2_ fold changes of commonly differentially expressed genes across different tissues. Log_2_ fold changes were calculated with methyl jasmonate-treated samples, comparing to control samples within each tissue type. Red represents upregulation, while blue represents downregulation. Tissue (in columns) and gene (in rows) clusters were calculated using default method in pheatmap to facilitate pattern identification. CG = cardiac glycoside Gene identifications are listed in Supplemental Table S3.

Methyl jasmonate treatment elicited expression changes in genes that are potentially involved in cardiac glycoside biosynthesis (Figure 8 and Table S3). Among those genes, two were annotated as *SMO* (squalene monooxygenase) and *SS1* (squalene synthase 1), which are involved in the final steps of terpenoid backbone biosynthesis (Kreis et al., 1998; Kreis and Müller-Uri, 2013; Zheng et al., 2014; Pandey et al., 2016; Zhang et al., 2017). One gene was annotated as *DWF1* or *DHCR24* (delta(24)-sterol reductase), which is associated with sterol biosynthesis to generate cholesterol for cardiac glycoside biosynthesis (Zerenturk et al., 2013; Tsukagoshi et al., 2016; Knoch et al., 2018).

We found induced expression of 14 cytochrome P450 genes from families that are involved in specialized metabolite biosynthesis in other species, including *CYP711A1, CYP714A1, CYP72A219, CYP81E8, CYP90B1, CYP94C1, CYP98A2*, *CYP450-AOS*, and *CYP450-HPL* (Figure 8 and Table S3). Whereas homologs of *CYP94C1, CYP450-AOS* and *CYP450-HPL* were annotated as being involved in jasmonic acid biosynthesis and signaling (Ruan et al., 2019; Li et al., 2022), other cytochrome P450s are potentially involved in cardiac glycoside biosynthesis.

In *D. lanata, D. purpurea*, *C. procera*, and *E. cheiranthoides*, known enzymes in the CYP87A family catalyze the predicted first step of cardiac glycoside biosynthesis, the conversion of campesterol to pregnenolone (Figure 9A) (Carroll et al., 2023; Kunert et al., 2023; Younkin et al., 2024). As *Asclepias* and *Calotropis* are both in the Apocynaceae, the *A. curassavica* enzyme catalyzing this reaction is likely to be similar to that of *C. procera.* Protein sequence comparisons and a phylogenetic analysis show that the *A. curassavica* AC04g009170.1 gene encodes a protein with 91% amino sequence identity to the cytochrome P450 that catalyzes the first step of cardiac glycoside biosynthesis in *C. procera* (Figures 9B and S5). This gene exhibited differential expression among *A. curassavica* tissue types, with highest expression in the stems and roots, and almost undetectable expression in the leaves, young leaves, buds, and flowers (Figure 9C). In the stems, there was a significant increase in *CYP87A* gene expression in response to methyl jasmonate treatment.

**Figure 9.**
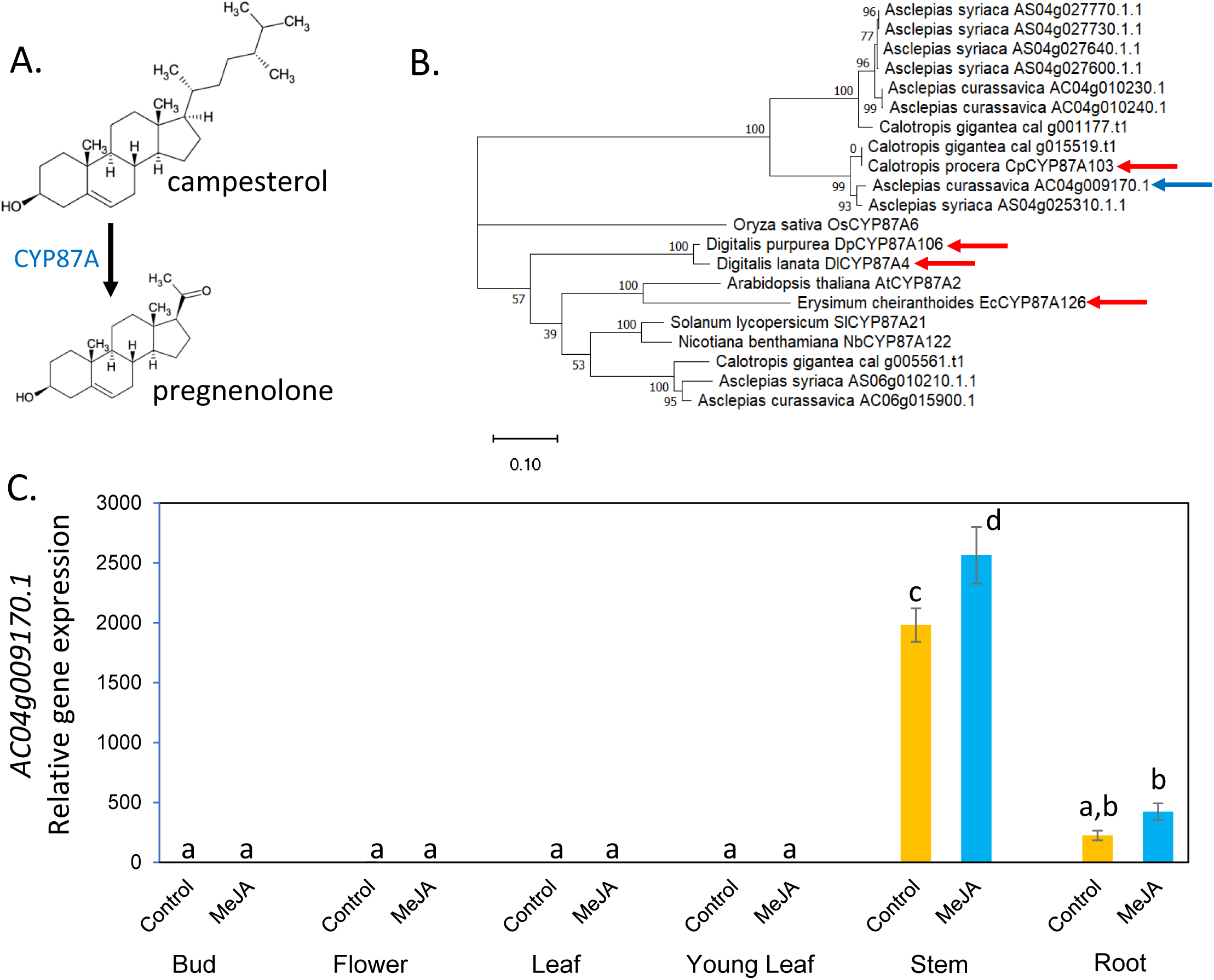
*Asclepias curassavica* CYP87A gene identification. A. First reaction of the cardiac glycoside biosynthesis pathway from campesterol, as described for *Digitalis* spp., *Erysimum cheiranthoides*, and *Calotropis procera*. B. Phylogenetic tree of predicted CYP87A proteins from cardiac glycoside-producing species, *Asclepias curassavica, Asclepias syriaca, C. procera, C. gigantea,* and *E. cheiranthoides*, as well as the well-studied model plant species *Nicotiana benthamiana, Solanum lycopersicum, Arabidopsis thaliana,* and *Oryza sativa.* Enzymes with confirmed functions in cardiac glycoside biosynthesis are marked with red arrows. The predicted *A. curassavica* enzyme catalyzing this reaction is marked with a blue arrow. The maximum likelihood, midpoint-rooted tree was produced with IQ-Tree and visualized with MEGA11. Bootstrap values are based on 1000 replicates. The scale bar indicates substitutions per site. A protein sequence alignment corresponding to this tree is presented in Figure S5. C. Expression of an *A. curassavica* CYP87A gene (AC04g009170.1, marked with a blue arrow in Panel B) in different tissue types, with and without methyl jasmonate (MeJA) treatment to elicit defense-related gene expression. Mean +/- s.e. of N = 8, different letters indicate P < 0.05, ANOVA followed by Tukey’s HSD test.

Beyond enzymes involved in the biosynthesis of the steroid core, modifying enzymes can generate cardiac glycoside diversity. For example, a permissive glycosyltransferase, UGT74AN1, catalyzes steroid C3 glycosylation to produce diverse glycosides and other bioactive molecules in *A. curassavica* (Wen et al., 2018). Our differential expression analysis revealed different groups of enzymes that may contribute to secondary modifications that generate the extensive cardiac glycoside diversity found in milkweed (Figure 8 and Table S3).

These enzymatic activities encoded by these genes include kinase, synthase, acyltransferase, glycosyl/glucosyltransferase, reductase, oxidase, dehydrogenase, ligase, hydrogenase, phosphatase, epimerase, esterase, cyclase, and UDPG glucosyltransferase. However, while confirming homology between our candidates to published sequences is possible, their functions in milkweed remain to be tested.

## Discussion

We have assembled and annotated a high-quality *A. curassavica* reference genome for use in ecological and molecular studies. By also sequencing the transcriptomes of six tissue types, with and without methyl jasmonate elicitation, we have identified both tissue-specific gene expression (Figure 7) and induced expression of genes that are likely to be involved in plant defenses (Figure 8, Figure S4, and Table S3). Previous milkweed research demonstrated tissue-specific accumulation of cardiac glycosides and resistance to insect herbivory (López-Goldar et al., 2022; Agrawal and Hastings, 2023). The tissue-specific milkweed transcriptomes described here will facilitate future research to identify the genetic basis of this within-plant natural variation in milkweed defenses. Additionally, correlation of gene expression and abundance of cardiac glycosides, *e.g*., voruscharin (Figure 1B), in different tissue types, may lead to identification of genes involved in the biosynthesis of these compounds.

Experiments with *D. lanata, D. purpurea*, *C. procera*, and *E. cheiranthoides* (Carroll et al., 2023; Kunert et al., 2023; Younkin et al., 2024) identified enzymes that catalyze conversion of campesterol to pregnenolone (Figure 9A) as the first step of cardiac glycoside biosynthesis. The *A. curassavica* genome encodes a CYP87A protein with 91% amino acid sequence identity to the confirmed enzyme from *C. procera* (Figures 9B and S5). The corresponding *A. curassavica* gene is expressed primarily in the roots and stems (Figure 9C). Thus, the initial step of *A. curassavica* cardiac glycoside biosynthesis may occur in stems and roots, which is different from what has been observed in both *Erysimum* and *Digitalis.* In *E. cheiranthoides*, cardiac glycosides are produced in the leaves and are transported to roots, flowers, and seeds (Alani et al., 2021). In *Digitalis*, cardiac glycoside biosynthesis occurs in the leaves, with mesophyll cells having the highest enzymatic activity (Hagimori et al., 1984). Although cardiac glycosides are not synthesized *de novo* in *Digitalis* roots (Hirotani and Furuya, 1977), root cells nevertheless can modify cardiac glycosides by hydroxylation, acetylation, and glycosylation (Alfermann et al., 1977).

Although the *A. curassavica* genome is smaller and has fewer LTR elements than the previously assembled *A. syriaca* genome (Figure 2), the two genomes are mostly colinear (Figure 3). Therefore, it is likely that results from functional genomic analyses with *A. curassavica* can be directly applied for research with *A. syriaca* and possibly other milkweed species for which genome sequences are not yet available. Comparative genomics studies will make it possible to identify genes that are present in one species but not another, *e.g.*, genes that are required for species-specific differences in the production of cardiac glycosides and other defense-related metabolites.

Prior to sequencing the *A. curassavica* genome, we conducted five generations of inbreeding by hand-pollinating individual plants. The resulting low level of DNA sequence heterozygosity facilitated our *A. curassavica* genome assembly. The availability of this inbred line will increase the reproducibility of future experiments to study the function of individual milkweed genes in ecological interactions and the biosynthesis of medically relevant specialized metabolites. To make our genome-sequenced *A. curassavica* lineage a permanent resource for milkweed researchers, we have submitted sixth-generation inbred seeds to a public repository, the USDA-ARS Ornamental Plant Germplasm Center (https://opgc.osu.edu/ ; https://www.ars-grin.gov/; accession number OPGC 7586).

### Data Availability

DNA and RNA sequences produced for this research have been deposited in GenBank, (PRJNA1055959). Seeds of the inbred line of *A. curassavica* that was used for genome sequencing are available from the USDA-ARS Ornamental Plant Germplasm Center, accession number OPGC 7586.

## Supplementary Materials

**Figure S1.** Genome heterozygosity, repeat content, and size estimate

**Figure S2.** Hi-C contact heat map of assembled pseudomolecules

**Figure S3.** Feature density along the *A. curassavica* pseudomolecules

**Figure S4.** Heat map of genes that are induced by methyl jasmonate across six tissue types

**Figure S5.** Alignment of CYP87A proteins

**Table S1.** Genome assembly metrics

**Table S2.** Analysis of *A. curassavica* gene families

**Table S3.** Differentially expressed genes

**Dataset S1.** Gene expression data for six *Asclepias curassavica* tissues, without methyl jasmonate treatment (control samples)

**Dataset S2.** Gene expression data for six *Asclepias curassavica* tissues, with methyl jasmonate treatment

**Dataset S3.** FASTA file of the protein sequences included in the Figure 9B phylogeny

**Dataset S4.** Machine-readable (Newick Format) file of the phylogenetic tree in Figure 9B

## Acknowledgements

This study was funded by US National Science Foundation awards 1645256 and 1850796, United States Department of Agriculture – National Institute of Food and Agriculture award 2020-67013-30896, and the Triad Foundation. We thank Arielle Johnson for assistance with the DNA sample preparation.

## Authors’ Contributions

SBB, FW, and GJ created an inbred line of *A. curassavica*; HF, AFP, JZ, LY, EY, and SRS assembled and annotated the genome; GLB conducted gene family analysis, HF generated and analyzed transcriptome data; and HF, GJ, and SRS wrote the manuscript.

