## Supplemental Figures S1-S5 for "Genome and tissue-specific transcriptome of the tropical milkweed (*Asclepias curassavica*)"

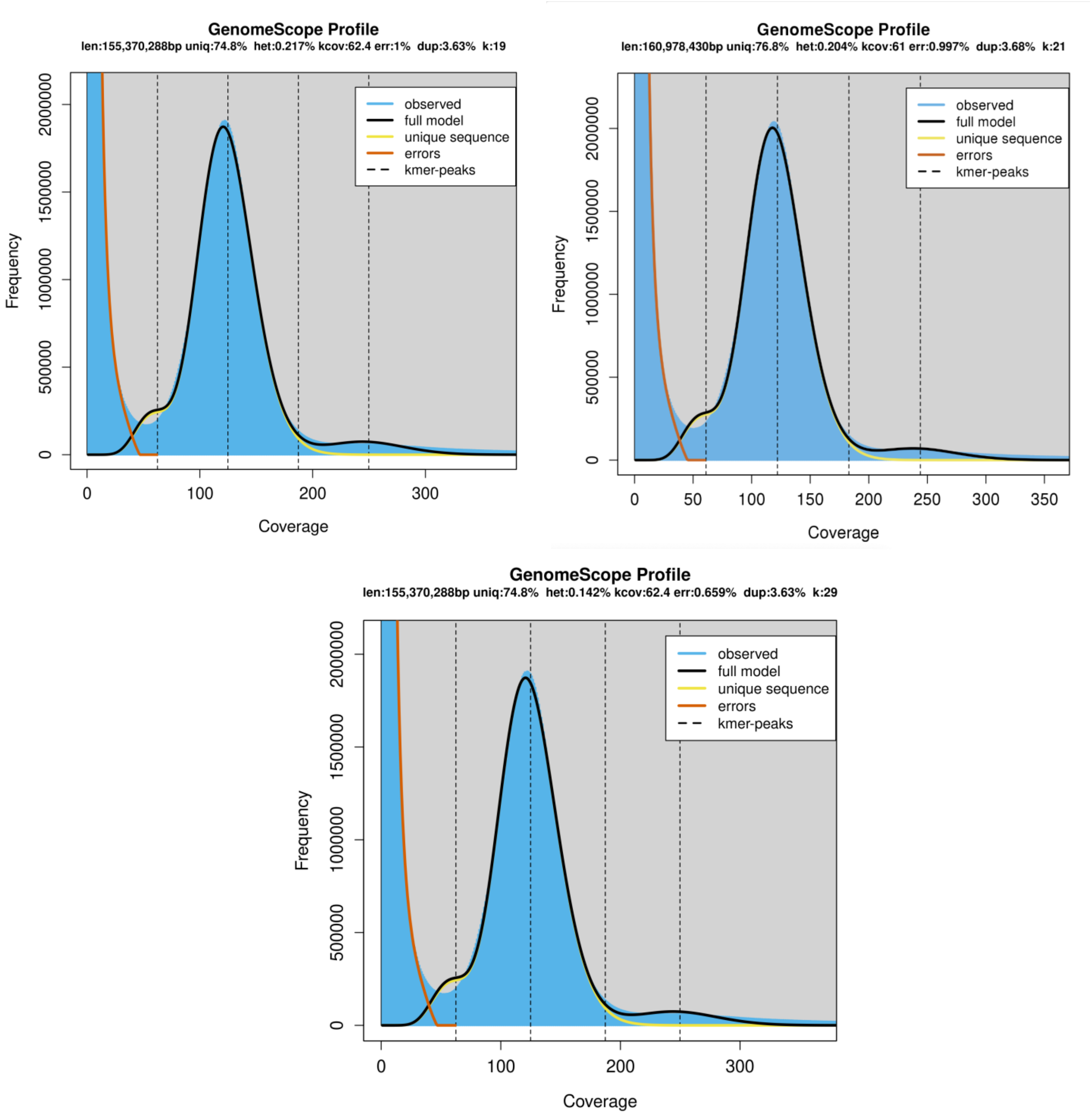


**Figure S1:** ***Asclepias curassavica*** **genome heterozygosity, repeat content, and size estimate based on K-mer analysis in GenomeScope.** Results were obtained from paired-end 2×150 bp sequencing reads. Plots for Kmer19, Kmer21, and Kmer29 are shown. Genome size estimates were not affected by Kmer length.

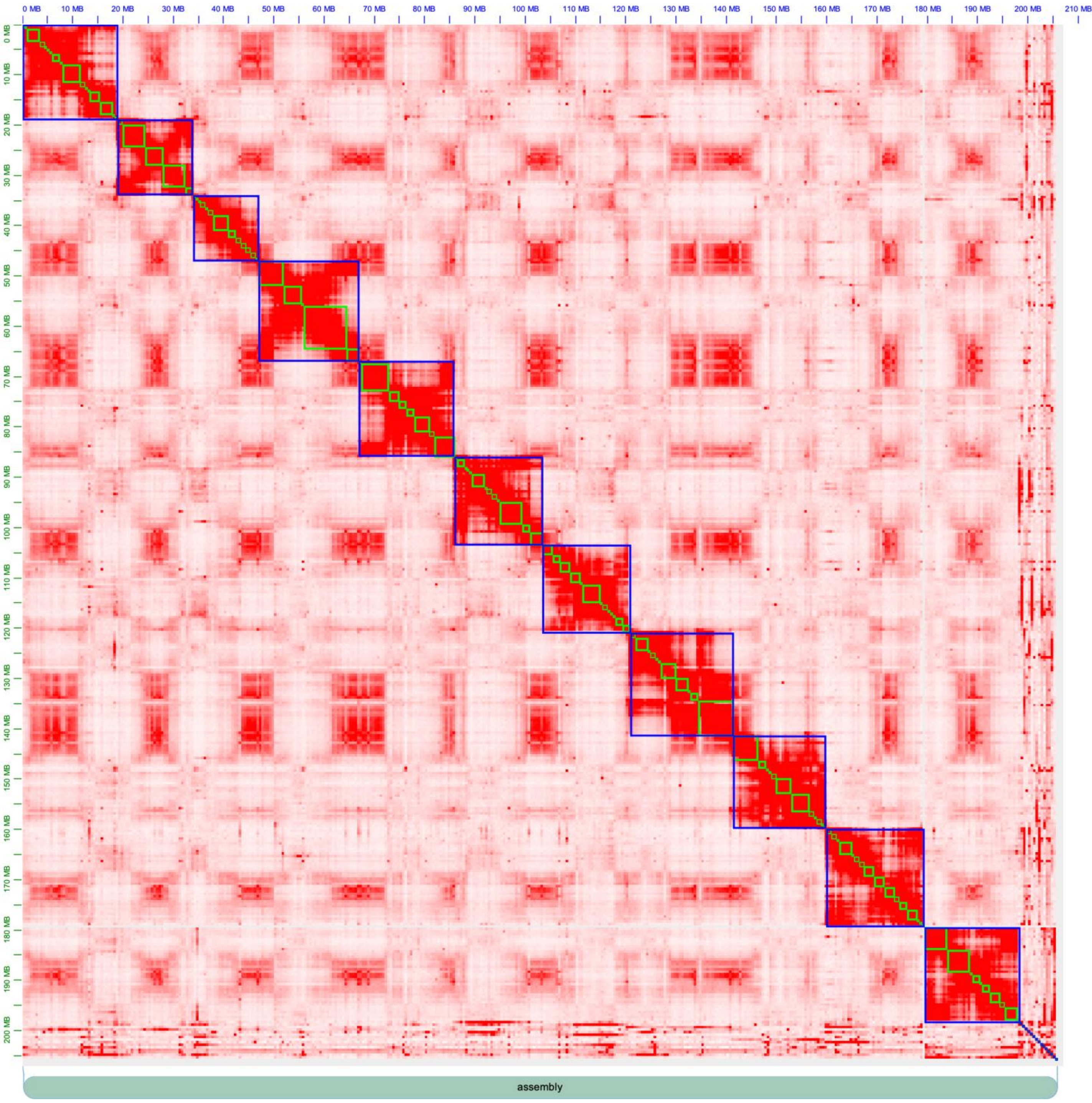


**FigureS2: Hi-C contact heat map of assembled pseudomolecules.** The largest scaffolds represent the 11 chromosomes of the *A. curassavica* genome.

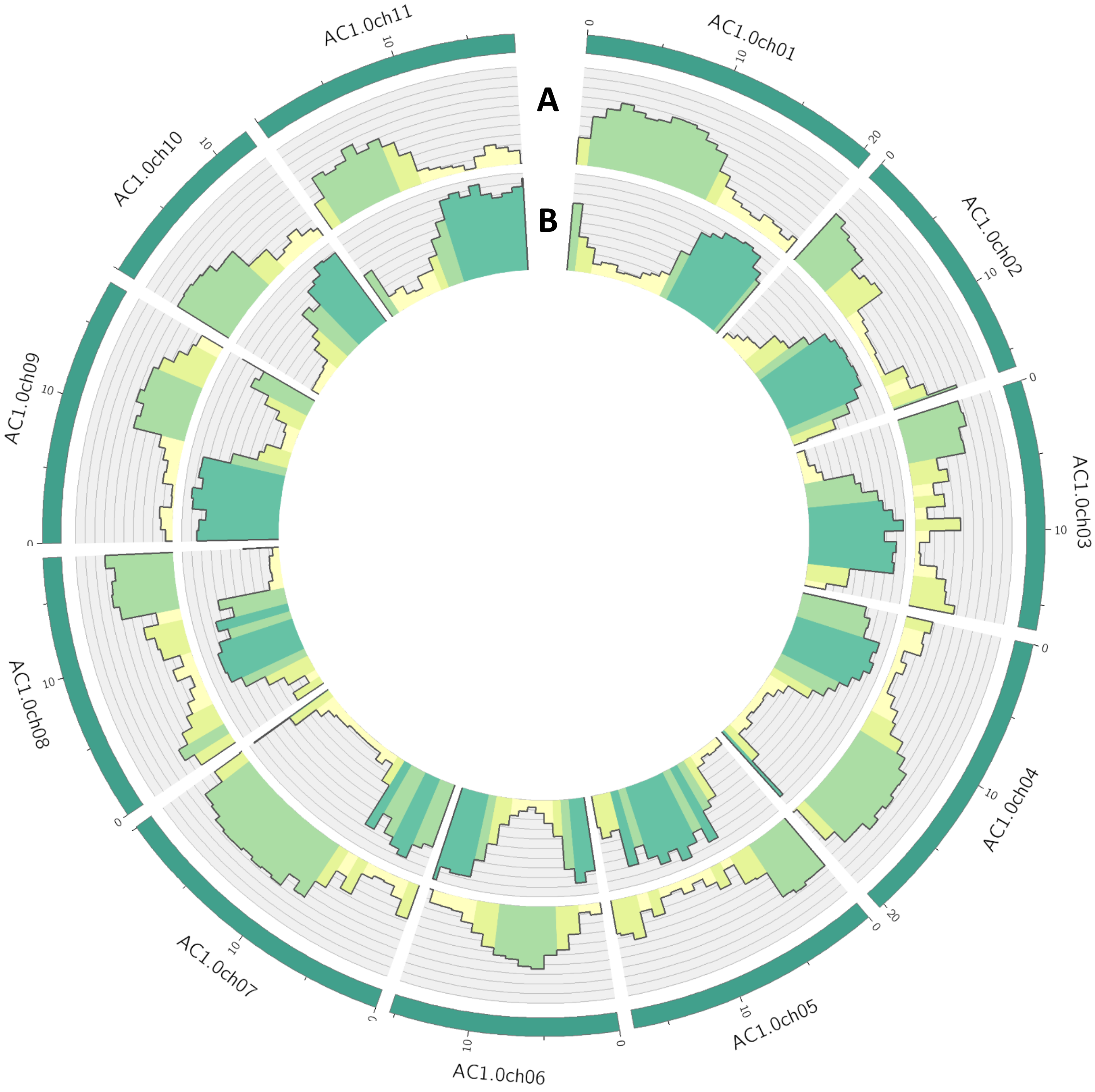


**Figure S3: Feature density along the *Asclepias curassavica* pseudomolecules.** Track A = gene density, track B = repeat density. Darker shading represents regions with greater feature density.

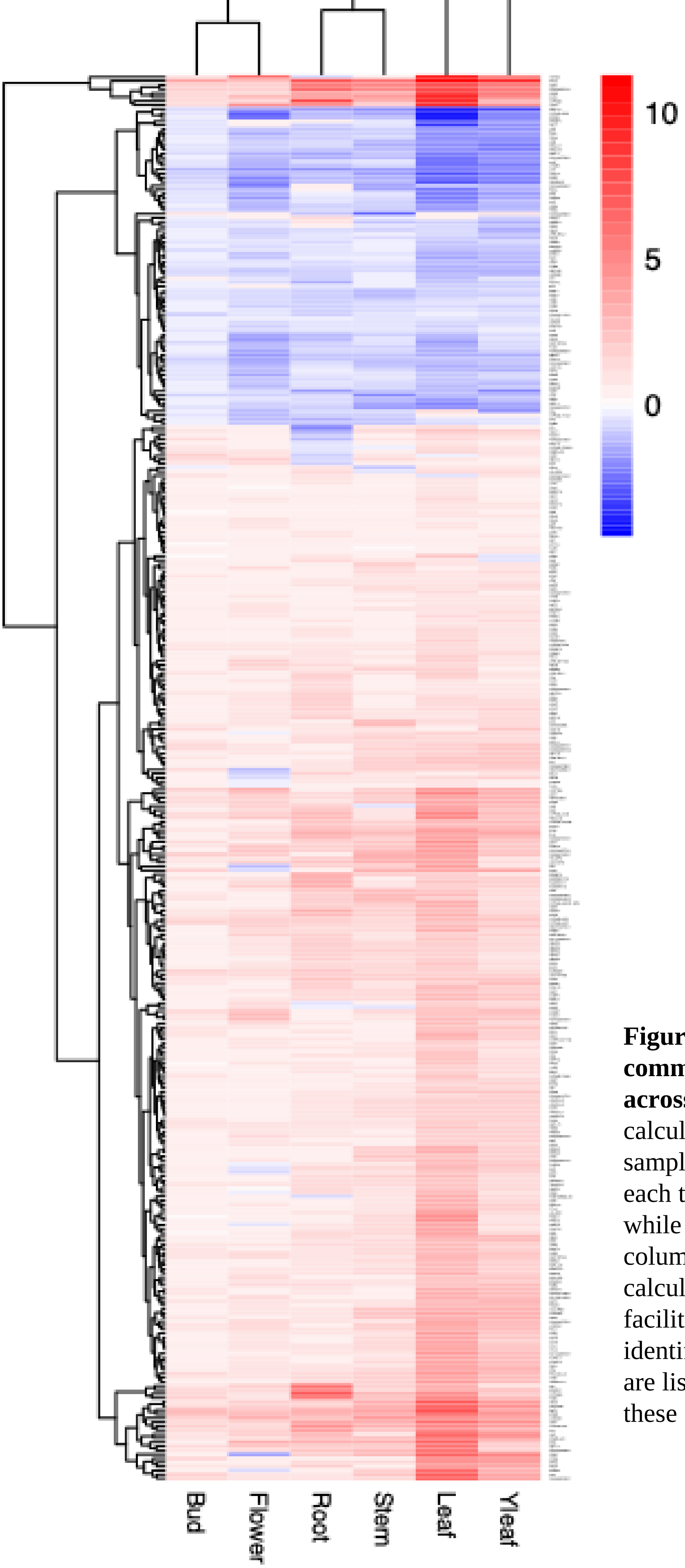


**Figure S4. Heatmap of $log_2$ fold changes of commonly differentially expressed genes across different tissues.** $Log_2$ fold changes were calculated with methyl jasmonate-treated samples, comparing to control samples within each tissue type. Red represents upregulation, while blue represents downregulation. Tissue (in columns) and gene (in rows) clusters were calculated using default method in pheatmap to facilitate pattern identification. Gene identifications and numerical expression values are listed in Supplemental Table S3. A subset of these genes are also shown in Figure 8.

```
Calotropis_gigantea_cal_g015519.t1        -----------------------------------------MMFAAIFLTTFLFIIVSRW     19
Calotropis_procera_CpCYP87A103            -----------------------------------------MMFAAIFLTTFLFIIVSRW     19
Asclepias_curassavica_AC04g009170.1       -----------------------------------------MMLAAILLTTFLFIILTRW     19
Asclepias_syriaca_AS04g025310.1.1         -----------------------------------------MMFAAIFLTTFLFIILSRW     19
Calotropis_gigantea_cal_g001177.t1        ------------------------------------------MVLAIALATILVILISRW     18
Asclepias_curassavica_AC04g010230.1       ---------------------------------KPSSNTYIKMVFAIALATILVILISRW     27
Asclepias_curassavica_AC04g010240.1       ---------------------------------KPSSNTYIKMVFAIALATILVILISRW     27
Asclepias_syriaca_AS04g027770.1.1         ------------------------------------------MVFAIALATILVILISRW     18
Asclepias_syriaca_AS04g027730.1.1         ------------------------------------------MVFAIALATILVILISRW     18
Asclepias_syriaca_AS04g027600.1.1         ------------------------------------------MVFAIALATILVILISRW     18
Asclepias_syriaca_AS04g027640.1.1         ------------------------------------------MVFAIALATILVILISRW     18
Oryza_sativa_OsCYP87A6                    MQPYLQLASLRLATTIPLAPRLYDANLLAASGAAMASSMAYIALLCA--ALAAVVALLRW     58
Digitalis_purpurea_DpCYP87A106            ---------------------------------------MSLVAISIGAILI--VIITNC     19
Digitalis_lanata_DlCYP87A4                ---------------------------------------MSLVAMSVGAILIIITIITNL     21
Calotropis_gigantea_cal_g005561.t1        ----------------------------------------MEVPVALCI-A-ALIIISITHW     20
Asclepias_syriaca_AS06g010210.1.1         ---------------------------FTS----L-GKMEVSVALCI-A-ALIIISITHW     26
Asclepias_curassavica_AC06g015900.1       ---------------------------FTS----L-SKMEVSVALCI-A-ALIIISITHW     26
Solanum_lycopersicum_SlCYP87A21           ---------------------------------------MISVGMSI-G-AFLILIIIHW     19
Nicotiana_benthamiana_NbCYP87A122         ---------------------------------------MISVGMCI-G-AFLVLIIIHW     19
Arabidopsis_thaliana_AtCYP87A2            -----------------------------------------MWALLIWV-SLLLISITHW     18
Erysimum_cheiranthoides_EcCYP87A126       ---------------------------------------MSWALCIWV-SLVVTGITTL     19
                                                                                          :

Calotropis_gigantea_cal_g015519.t1        IYRWRNPS--CNGILPPGSMGLPIIGESLAYFTPYFKDDIPLFVRERVQKYGPLFRTSLV     77
Calotropis_procera_CpCYP87A103            IYRWRNPS--CNGILPPGSMGLPIIGESLAYFTPYFKDDIPLFVRERVQKYGPLFRTSLV     77
Asclepias_curassavica_AC04g009170.1       IYRWRNPS--CNGILPPGSMGLPIIGESLAYFNPYFKDDIPLFVRKRVQKYGPLFRTSLV     77
Asclepias_syriaca_AS04g025310.1.1         IYRWRNPS--CNGILPPGSMGLPIIGESLAYFTPYFKDDIPAFVRKRVQKYGPLFRTSLV     77
Calotropis_gigantea_cal_g001177.t1        VYKWRNPS--CNGVLPPGSMGLPIIGESIAYFTPYFKDDVPLFIRKRVAKYGPLFRTSIV     76
Asclepias_curassavica_AC04g010230.1       VYKWRNPS--CNGVLPPGSMGLPIIGESIAYFTPYFKDDVPLFIRKRVAKYGPLFRTSIV     85
Asclepias_curassavica_AC04g010240.1       VYKWRNPS--SNGVLPPGSMGLPIIGESIAYFTPYFKDDVPLFIRKRVAKYGPLFRTSIV     85
Asclepias_syriaca_AS04g027770.1.1         VYKWRNPS--CNGVLPPGSMGLPIIGESIAYFTPYFKDDVPLFIRKRVAKYGPLFRTSIV     76
Asclepias_syriaca_AS04g027730.1.1         VYKWRNPS--CNGVLPPGSMGLPIIGESIAYFTPYFKDDVPLFIRKRVAKYGPLFRTSIV     76
Asclepias_syriaca_AS04g027600.1.1         VYKWRNPS--CNGVLPPGSMGLPIIGESIAYFTPYFKDDVPLFIRKRVAKYGPLFRTSIV     76
Asclepias_syriaca_AS04g027640.1.1         VYKWRNPS--CNGVLPPGSMGLPIIGESIAYFTPYFKDDVPLFIRKRVAKYGPLFRTSIV     76
Oryza_sativa_OsCYP87A6                    AYRWSHPR--SNGRLPPGSLGLPVIGETLQFFAPNPTCDLSPFVKERIKRYGSIFKTSVV     116
Digitalis_purpurea_DpCYP87A106            VFKWRNRSLSSGGILPPGSFGWPLIGETLHFFTPNTSFDVTPFVKDRMKRYGPIFKTSLV     79
Digitalis_lanata_DlCYP87A4                VFKWWNRSLSSGGVLPPGSFGWPLIGETLHFFTPNTSFDVTPFVKDRMKRYGPIFKTSLV     81
Calotropis_gigantea_cal_g005561.t1        IYRWKNPK--CKGKLPPGSMGWPLLGETLPFFAPTSTFDVHPFVKERMQRYGPIFRTSLV     78
Asclepias_syriaca_AS06g010210.1.1         IYRWKNPK--CNGKLPPGSMGWPLLGETLPFFAPSSSFDVHPFVKERMQRYGPIFRTSLV     84
Asclepias_curassavica_AC06g015900.1       IYRWKNPK--CNGKLPPGSMGWPLLGETLPFFAPSSSFDVHPFVKERMQRYGPIFRTSLV     84
Solanum_lycopersicum_SlCYP87A21           VYNWRNPR--CNGKLPPGSMGWPLLGETIQFFTPNTTLDIAPFVKERMKRYGPIFRTSVV     77
Nicotiana_benthamiana_NbCYP87A122         VYNWRNPR--CNGKLPPGSMGWPLLGETIPFFAPNTSSDIAPFVKDRMQRYGPIFRTSVV     77
Arabidopsis_thaliana_AtCYP87A2            VYSWRNPK--CRGKLPPGSMGFPLLGESIQFFKPNKTSDIPPFIKERVKKYGPIFKTNLV     76
Erysimum_cheiranthoides_EcCYP87A126       VYKWRNPK--CSGKLPPGSMGLPLLGETIQFFKPNLTSDIQPFIKERTKKYGPIFKTSLV     77
                                           : * :    . * *****:* *::**:: :* *  . *:  *::.*  :** :*:*.:*

Calotropis_gigantea_cal_g015519.t1        GQSVIVSTDPEVNYYIFQQEGNLFQCWYSESVLKVLGEQSMAVQAGAFHKYLKNLCLSLV     137
Calotropis_procera_CpCYP87A103            GQSVIVSTDPEVNYYIFQQEGNLFQCWYSESVLKVLGEQSMAVQAGAFHKYLKNLCLSLV     137
Asclepias_curassavica_AC04g009170.1       GQSVIVSTDPEVNYYIFQQEGNLFQCWYSESVLKVLGEQSMAVQAGAFHKYLKNLCLSLV     137
Asclepias_syriaca_AS04g025310.1.1         GQSVIVSTDPEVNYYIFQQEGNLFQCWYSESVLKVLGEQSMAVQAGAFHKYLKNLCLSLV     137
Calotropis_gigantea_cal_g001177.t1        GQPVVVSTDPEVNYYVFQQEGNIFQCWYFESVNRIIGEQSMAVQQGVVHKFLKNLILTLV     136
Asclepias_curassavica_AC04g010230.1       GQPVVVSTDPEVNYYVFQQEGNIFQCWYFESVNRIIGQQSMAVQQGVVHKYLKNLILSLV     145
Asclepias_curassavica_AC04g010240.1       GQPVVVSTDPEVNYYVFQQEGNIFQCWYFESVNRIIGQQSMAVQQGVVHKYLKNLILSLV     145
Asclepias_syriaca_AS04g027770.1.1         GQPVVVSTDPEVNYYVFQQEGNIFQCWYFESVNRIIGQQSMAVQQGVVHKYLKNLILSLV     136
Asclepias_syriaca_AS04g027730.1.1         GQPVVVSTDPEVNYYVFQQEGNIFQCWYFESVNRIIGQQSMAVQQGVVHKYLKNLILSLV     136
Asclepias_syriaca_AS04g027600.1.1         GQPVVVSTDPEVNYYVFQQEGNIFQCWYFESVNRIIGQQSMAVQQGVVHKYLKNLILSLV     136
Asclepias_syriaca_AS04g027640.1.1         GQPVVVSTDPEVNYYVFQQEGNIFQCWYFESVNRIIGQQSMAVQQGVVHKYLKNLILSLV     136
Oryza_sativa_OsCYP87A6                    GRPVVVSADPEMNYYVFQQEGKLFESWYPDTFTEIFGRDNVGSLHGFMYKYLKTLVLRLY     176
Digitalis_purpurea_DpCYP87A106            GVPVIVSTDAELNNFIFQQEGQTFQSWYPSTFTEIFGRENLSTLHGFMYKYFKNMVLGLF     139
Digitalis_lanata_DlCYP87A4                GVPVIVSTDAELNNFIFQQEGQTFQSWYPSTFTEIFGRENLSTLHGFMYKYFKNMVLGLF     141
Calotropis_gigantea_cal_g005561.t1        GRPVIVSTDSDLNYFIFQQEGQLFQSWYPDTFTEIFGRQNVGSLHGFMYKYLKNMMLNLF     138
Asclepias_syriaca_AS06g010210.1.1         GRPVIVSTDSDLNYFIFQQEGQLFQSWYPDTFTEIFGRQNIGSLHGFMYKYLKNMMLNLF     144
Asclepias_curassavica_AC06g015900.1       GRPVIVSTDSDLNYFIFQQEGQLFQSWYPDTFTEIFGRQNIGSLHGFMYKYLKNMMLNLF     144
Solanum_lycopersicum_SlCYP87A21           GRPVIVSTDPDLNYFIFQQEGQSFQSWYPDTFTEIFGRQNVGSLHGFMYKYLKNMVLNLF     137
Nicotiana_benthamiana_NbCYP87A122         GRPVIASTDPDLNYFIFQQEGQLFQSWYPDTFTEIFGKQNVGSLHGFMYKYLKNMVLNLF     137
Arabidopsis_thaliana_AtCYP87A2            GRPVIVSTDADLSYFVFNQEGRCFQSWYPDTFTHIFGKKNVGSLHGFMYKYLKNMVLTLF     136
Erysimum_cheiranthoides_EcCYP87A126       GKSIIVTTDPDFSYFVFQQEGQSFQSWYPDTFVEIFGKQNLGALHGIIYKYLKHMVLSLF     137
                                          *  ::.::* :.. ::*:***. *:.** .:. .::*...:.   * .:*::* : * *
```

```
Calotropis_gigantea_cal_g015519.t1      GPENLKETLMYEMDQNTIEHLQSWGT-IGNLDAKDATAELVFKLAARKIINYDEKKS-GK    195
Calotropis_procera_CpCYP87A103          GPENLKETLMYEMDQNTIEHLQSWGT-IGNLDAKDATAELVFKLAARKIINYDEKKS-GK    195
Asclepias_curassavica_AC04g009170.1     GPENLKETLMHEMDQNTTQHLLSWGT-IGNLDAKDATAELVFKLAAKKILNYDEKKS-GK    195
Asclepias_syriaca_AS04g025310.1.1       GPESLKETLMHEMDQNTIQHLLSWGT-IGNLDAKDATAELVFKLAAKKILNYDEKKS-GK    195
Calotropis_gigantea_cal_g001177.t1      GPENLKEKLILEMDQNTRQYLHSWAN-IGNIDAKDATAEMVFTLAAKKILNYDDKKA-SK    194
Asclepias_curassavica_AC04g010230.1     GPENLKEKLILEMDQNTRQYLQSWAN-MGNLDAKDATAEMVFTLAAKKILNYDDKKA-SK    203
Asclepias_curassavica_AC04g010240.1     GPENLKEKLILEMDQNTRQYLQSWAN-MGNLDAKDATAEMVFTLAAKKILNYDDKKA-SK    203
Asclepias_syriaca_AS04g027770.1.1       GPENLKEKLILEMDQNTRQYLQSWAN-MGNLDAKDATAEMVFTLAAKKILNYDDKKA-SK    194
Asclepias_syriaca_AS04g027730.1.1       GPENLKEKLILEMDQNTRQYLQSWAN-MGNLDAKDATAEMVFTLAAKKILNYDDKKA-SK    194
Asclepias_syriaca_AS04g027600.1.1       GPENLKEKLILEMDQNTRQYLQSWAN-MGNLDAKDATAEMVFTLAAKKILNYDDKKA-SK    194
Asclepias_syriaca_AS04g027640.1.1       GPENLKEKLILEMDQNTRQYLQSWAN-MGNLDAKDATAEMVFTLAAKKILNYDDKKA-SK    194
Oryza_sativa_OsCYP87A6                  GQENLKSVLLAETDAACRGSLASWAS-QPSVELKEGISTMIFDLTAKKLIGYDPSKPSQV    235
Digitalis_purpurea_DpCYP87A106          GPESLKTMISEVENT-SNINLKRWSA-NGTVELKDAIAEMIFELTAKKLISYELEKS-PY    196
Digitalis_lanata_DlCYP87A4              GPESLKTMISEVENT-SNINLKRWSSSNGTVELKDAIAEMIFELTAKKLISYELEKS-PY    199
Calotropis_gigantea_cal_g005561.t1      GPEALKKMIPEVEQV-AKRKLREWSS-QTTTEMKEATASMIFDLTAKKLISYDSEKS-SD    195
Asclepias_syriaca_AS06g010210.1.1       GPEALKKMIPEVEQV-AKRNLRKWSS-QTTTEMKEATASMIFHLTAKKLISYDSEKS-SD    201
Asclepias_curassavica_AC06g015900.1     GPEALKKMIPEVEQV-AKRNLRKWSS-QTTTEMKEATASMIFHLTAKKLISYDSEKS-SD    201
Solanum_lycopersicum_SlCYP87A21         GSESLKKMLPEVEEV-AKNKLKRWSG-QTSVEMKEATANMIFDLTAKKLISYDSETS-SE    194
Nicotiana_benthamiana_NbCYP87A122       GPESLKKMMPEVEEA-AKNKLKRWSG-QTSVEMKEATANMIFDLTAKKLISYDSENS-SE    194
Arabidopsis_thaliana_AtCYP87A2          GHDGLKKMLPQVEMT-ANKRLELWSN-QDSVELKDATASMIFDLTAKKLISHDPDKS-SE    193
Erysimum_cheiranthoides_EcCYP87A126     GFESLKNMLPEIEQT-ACKKLDLWST-QKSIELKESTANLIFDLTAKKLISHDEEKS-SE    194
                                        * : **  :            *  *.    . : *:. : ::* *:*:*::.:: ..

Calotropis_gigantea_cal_g015519.t1      KLRDCYKAFMDGFISFPLYIPGTAFYACIQ---------------GRKKALKVIKEVFNQ    240
Calotropis_procera_CpCYP87A103          KLRDCYKAFMDGFISFPLYIPGTAFYACIQ---------------GRKKALKVIKEVFNQ    240
Asclepias_curassavica_AC04g009170.1     KLRDCYKAFMDGFISFPLYIPGTAFYACIQ------------------------------    225
Asclepias_syriaca_AS04g025310.1.1       KLRDCYKAFMDGFISFPLYIPGTAFYACIQ---------------GRKRALKLIKEIFNQ    240
Calotropis_gigantea_cal_g001177.t1      ELRDCYKAFLDGFISFPLYIPGTAFYACIQ---------------GRRKALKVIKNIFNE    239
Asclepias_curassavica_AC04g010230.1     ELRDCYKAFLDGFISFPLYIPGTAFYACIQ---------------GRRKALKVIKNIFNE    248
Asclepias_curassavica_AC04g010240.1     ELRDCYKAFLDGFISFPLYIPGTAFYACIQ---------------GRRKALKVIKNIFNE    248
Asclepias_syriaca_AS04g027770.1.1       ELRDCYKAFLDGFISFPLYIPGTAFYACIQ---------------GRRKALKVIKNIFNE    239
Asclepias_syriaca_AS04g027730.1.1       ELRDCYKAFLDGFISFPLYIPGTAFYACIQ---------------GRRKALKVIKNIFNE    239
Asclepias_syriaca_AS04g027600.1.1       ELRDCYKAFLDGFISFPLYIPGTAFYACIQ---------------GRRKALKVIKNIFNE    239
Asclepias_syriaca_AS04g027640.1.1       ELRDCYKAFLDGFISFPLYIPGTAFYACIQ---------------GRRKALKVIKNIFNE    239
Oryza_sativa_OsCYP87A6                  NLRKNFGAFICGLISFPLNIPGTAYHECME---------------GRKNAMKVLRGMMKE    280
Digitalis_purpurea_DpCYP87A106          NLRDNFVAFIDGLISFPLNIPGTAYYKCLQ---------------GRKNAIKMLRDMLHE    241
Digitalis_lanata_DlCYP87A4              NLRDNFVAFIDGLISFPLNIPGTAYYRCLQ---------------GRKNAIKMLKDMLHE    244
Calotropis_gigantea_cal_g005561.t1      NLRESFVAFMQGLISFPLDIPGTAYHQCMQ---------------GRKKAMKMLTNMLNE    240
Asclepias_syriaca_AS06g010210.1.1       NLRESFVAFIQGLISFPLDIPGTAYHQCMQPLPPSLPPSLSQEPGGRKKAMKMLTNMLNE    261
Asclepias_curassavica_AC06g015900.1     NLRESFVAFIQGLISFPLDIPGTAYHQCMQGILYIIDDSSCLIKQGRKKAMKMLTNMLNE    261
Solanum_lycopersicum_SlCYP87A21         NLRESFVAFIQGLISFPIDIPGTAYHKCLQ---------------GRKKAMKMLKTMLEE    239
Nicotiana_benthamiana_NbCYP87A122       NLRESFVAFIQGLISFPIDIPGTVYHKCLQ---------------GRKKAMKMLKTLLEE    239
Arabidopsis_thaliana_AtCYP87A2          NLRANFVAFIQGLISFPFDIPGTAYHKCLQ---------------GRAKAMKMLRNMLQE    238
Erysimum_cheiranthoides_EcCYP87A126     NLRDNYVAFIDGLISFPINLPGTAFYKCLK---------------GRERVMSSLRNMLKE    239
                                        :**  : **: *:****: :***.:: *::

Calotropis_gigantea_cal_g015519.t1      RRGIGATE------EKQKVFVDYILEEVDNKESFITEGIALDLVFLLLFASHETTSTAMT    294
Calotropis_procera_CpCYP87A103          RRGIGATE------EKQKVFVDYILEEVDNKESFITEGIALDLVFLLLFASHETTSTAMT    294
Asclepias_curassavica_AC04g009170.1     -----------------KVFVDYVLEEVDNKESFITEGIAQDLVFLLLFASHETTSTAMT    268
Asclepias_syriaca_AS04g025310.1.1       RRGIGETE------EKQKVFVDYVLEEVDNKESFITEGIALDLVFLLLFASHETTSTAMT    294
Calotropis_gigantea_cal_g001177.t1      RRVSAST-SME---KKKNDFVDTVLEQVDSKDSFLNEEIALDLVFLLLFASHETTSTAMT    295
Asclepias_curassavica_AC04g010230.1     RRAIAGSNSMEKK---KNDFVDAVLEQVDSKDSFLNEEIALDLVFLLLFASHETTSTAMT    305
Asclepias_curassavica_AC04g010240.1     RRAIAGSNSMEKK---KNDFVDAVLEQVDSKDSFLNEEIALDLVFLLLFASHETTSTAMT    305
Asclepias_syriaca_AS04g027770.1.1       RRANANSNSMEKKKKEKNDFVDAVLEQVDSKDSFLSEEIALDLVFLLLFASHETTSTAMT    299
Asclepias_syriaca_AS04g027730.1.1       RRANANSNSMEKKKKEKNDFVDAVLEQVDSKDSFLSEEIALDLVFLLLFASHETTSTAMT    299
Asclepias_syriaca_AS04g027600.1.1       RRANA--NSMEKKKKEKNDFVDAVLEQVDSKDSFLSEEIALDLVFLLLFASHETTSTAMT    297
Asclepias_syriaca_AS04g027640.1.1       RRANA--NSMEKKKKEKNDFVDAVLEQVDSKDSFLSEEIALDLVFLLLFASHETTSTAMT    297
Oryza_sativa_OsCYP87A6                  RMAEPE--------RPCEDFFDHVIQELRREKPLLTETIALDLMFVLLFASFETTALALT    332
Digitalis_purpurea_DpCYP87A106          RREKPR--------ETQTDFFDYVLEELQKQDTIITETLALDLMFVLLFASHETASIALT    293
Digitalis_lanata_DlCYP87A4              RREKPR--------ETQTDFFDYVLEELQKEDTIITETLALDLMFVLLFASHETASIALT    296
Calotropis_gigantea_cal_g005561.t1      RRANPK--------KHSTDFFDFVLEELGRKNTILTEAIALDLMFVLLFASFETTSLALT    292
Asclepias_syriaca_AS06g010210.1.1       RRANPK--------KHSTDFFDFVLEELGRKNTILTEAIALDLMFVLLFAN---NFLGTN    310
Asclepias_curassavica_AC06g015900.1     RRANPK--------KHSTDFFDFVLEELGRKNTILTEAIALDLMFVLLFASFETTSLALT    313
Solanum_lycopersicum_SlCYP87A21         RRAKPR--------KEGTDFFDYVLEELQKNDIILTEAIALDLMFVLLFASFETTSLAIT    291
Nicotiana_benthamiana_NbCYP87A122       RRAKPR--------KEQSDFFDYVLEELQRKDTILTEAIALDLMFVLLFASFETTSLAIT    291
Arabidopsis_thaliana_AtCYP87A2          RRENPR--------KNPSDFFDYVIEEIQKEGTILTEEIALDLMFVLLFASFETTSLALT    290
Erysimum_cheiranthoides_EcCYP87A126     RRKNPR--------KVASDFFDYVIEELKKEGTMLTESIALDLMFVLLFASFETTSLAIT    291
                                                           *.* ::::  :  ::.* :* **:*:****.      . .
```

```
Calotropis_gigantea_cal_g015519.t1       MAMKFITESPAVLAELVREHEAILKNRENP---ESGITWKEYKGMTFTHMVINETVRIAN     351
Calotropis_procera_CpCYP87A103           MAMKFITESPAVLAELVREHEAILKNRENP---ESGITWKEYKGMTFTHMVINETVRIAN     351
Asclepias_curassavica_AC04g009170.1      MAMKFITESPAVLAELVREHEAILKNRENP---ESGITWKEYKGMTFTHMVINETVRIAN     325
Asclepias_syriaca_AS04g025310.1.1        MAMKFITESPAVLAELVREHEAILKNRENP---ESGITWKEYKGMTFTHMVINETVRIAN     351
Calotropis_gigantea_cal_g001177.t1       LAMKYLSQHPAVLAELL-----------------------------------VINETVRLAN     322
Asclepias_curassavica_AC04g010230.1      LAMKYLSLHPAVLAELLREHETILSNREGTETASSPITWKEYKSMTFTHMVINETVRLAN     365
Asclepias_curassavica_AC04g010240.1      LAMKYLSLHPAVLAELLREHETILSNREGTETASSPITWKEYKSMTFTHMVINETVRLAN     365
Asclepias_syriaca_AS04g027770.1.1        LAMKYLSLHPAVLAELLREHETILSNREGTETAASPITWKEYKSMTFTHMVINETVRLAN     359
Asclepias_syriaca_AS04g027730.1.1        LAMKYLSLHPAVLAELLREHETILSNREGTETAASPITWKEYKSMTFTHMVINETVRLAN     359
Asclepias_syriaca_AS04g027600.1.1        LAMKYLSLHPAVLAELLREHETILSNREGTETAASPITWKEYKSMTFTHMVINETVRLAN     357
Asclepias_syriaca_AS04g027640.1.1        LAMKYLSLHPAVLAELLREHETILSNREGTETAASPITWKEYKSMTFTHMVINETVRLAN     357
Oryza_sativa_OsCYP87A6                   IGVKLLTENPKVVDALREEHEAIIRNRKDP---NSGVTWAEYKSMTFTSQVIMEIVRLAN     389
Digitalis_purpurea_DpCYP87A106           LAMKFLVDHPLVLDKLTEEHDEIIKMREDP---NSGLTWNEYKSMKFTFQFINETLRLAN     350
Digitalis_lanata_DlCYP87A4               LAMKFLVDHPLVLEKLTEEHDEIIKTREDP---NSGLTWNEYKSMKFTFQFINETLRLAN     353
Calotropis_gigantea_cal_g005561.t1       LATKLLVDHPLALKALTEEHEAIIKKRENP---DSALTWSEYKSMTFTFQFINETVRMAN     349
Asclepias_syriaca_AS06g010210.1.1        LSNKLLVDHPLALKALTEEHEAIIKKRENP---DSALTWSEYKSMTFTFQFINETVRMAN     367
Asclepias_curassavica_AC06g015900.1      LATKLLVDHPLALKALTEEHETIIKNRENP---DSALTWSEYKSMTFTFQFINETVRMAN     370
Solanum_lycopersicum_SlCYP87A21          LATKFLHDHPLALKELTEEHEAIIRSRENP---ASGLTWKEYKSMKFTFQVINETVRLAN     348
Nicotiana_benthamiana_NbCYP87A122        LATKFLHDHPLALKELTEEHEAIIRRRENP---ASGLTWKEYKSMKFTFQVINETVRLAN     348
Arabidopsis_thaliana_AtCYP87A2           LAIKFLSDDPEVLKRLTEEHETILRNREDA---DSGLTWEEYKSMTYTFQFINETARLAN     347
Erysimum_cheiranthoides_EcCYP87A126      VAIKMLSDHPSVLKRLTEEHEAILRNRKDP---NSGLTWEEYKSMTYTFQFMNETARLAN     348
                                         :. * :   * .:  *                                  .: *  *:**

Calotropis_gigantea_cal_g015519.t1       IAPGIFRKVMKEVEIKGYTIPAGWTVMVCPSTVHMNPDKYENPLSFDPWRWEGQ-ELHSA     410
Calotropis_procera_CpCYP87A103           IAPGIFRKVMKEVEIKGYTIPAGWTVMVCPSTVHMNPDKYENPLSFDPWRWEGQ-ELHSA     410
Asclepias_curassavica_AC04g009170.1      IAPGIFRKVMKEVEIKGYTIPAGWTVMVCPSTVHMNPDKYENPLSFDPWRWEGQ-ELHSA     384
Asclepias_syriaca_AS04g025310.1.1        IAPGIFRKVMKEVEIKGYTIPAGWTVMVCPSTVHMNPDKYENPLSFDPWRWEGQ-ELHSA     410
Calotropis_gigantea_cal_g001177.t1       IAPGIFRKVMKEVEIK------GWTLMVCPSSVHMNADKYEDPLAFNPWRWEGQ-ELHSA     375
Asclepias_curassavica_AC04g010230.1      IAPGIFRKVMKEVEIKGYTIPEGWTLMVCPSSVHMNGDKYEDPLEFNPWRWEGQ-ELHSA     424
Asclepias_curassavica_AC04g010240.1      IAPGIFRKVMKEVEIKGYTIPEGWTLMVCPSSVHMNGDKYEDPLEFNPWRWEGQ-ELHSA     424
Asclepias_syriaca_AS04g027770.1.1        IAPGIFRKVMKEVEIKGYTIPEGWTLMVCPSSVHLNADKYEDPLEFNPWRWEGQ-ELHSA     418
Asclepias_syriaca_AS04g027730.1.1        IAPGIFRKVMKEVEIKGYTIPEGWTLMVCPSSVHLNADKYEDPLEFNPWRWEGQ-ELHSA     418
Asclepias_syriaca_AS04g027600.1.1        IAPGIFRKVMKEVEI-----KEGWTLMVCPSSVHMNADKYEDPLEFNPWRWEGQ-ELHSA     411
Asclepias_syriaca_AS04g027640.1.1        IAPGIFRKVMKEVEIKGYTIPEGWTLMVCPSSVHMNADKYEDPLEFNPWRWEGQ-ELHSA     416
Oryza_sativa_OsCYP87A6                   IVPGIFRKALQDVEIKGYTIPAGWGIMVCPPAVHLNPEIYEDPLAFNPWRWQGKPEITGG     449
Digitalis_purpurea_DpCYP87A106           IAPLIFRKALTETEFKGYTIPAGWAVMVCLPAVHLDPTKYKNPLEFNPWRWEGV-DTSVG     409
Digitalis_lanata_DlCYP87A4               IAPLIFRKALTETEFKGYTIPAGWAVMVCLPAVHLDPSKYKNPLEFNPWRWEGV-DTSVG     412
Calotropis_gigantea_cal_g005561.t1       IVPGIFRKALKDINFKGYTIPAGWAVMVCPPAVHMNPGKYQNPLEFNPWRWEGM-ELNGA     408
Asclepias_syriaca_AS06g010210.1.1        IVPGIFRKALKDINFKGYTIPAGWAVMVCPPAVHMNPGKYQNPLEFNPWRWEGM-ELNGA     426
Asclepias_curassavica_AC06g015900.1      IVPGIFRKALKDINFKGYTIPAGWAIMVCPPAVHMNPGKYQNPLEFNPWRWEGM-ELNGA     429
Solanum_lycopersicum_SlCYP87A21          IVPAIFRKTLTDINFKGYTIPAGWAVMVCPPAVHLNPAKYQDPLDFNPWRWEGV-EINGA     407
Nicotiana_benthamiana_NbCYP87A122        IVPAIFRKALRDVNFKGYTIPAGWAIMVCPPAVHLNPAKYQDPLEFNPWRWEGV-EMNGA     407
Arabidopsis_thaliana_AtCYP87A2           IVPAIFRKALRDIKFKDYTIPAGWAVMVCPPAVHLNPEMYKDPLVFNPSRWEGS-KVTNA     406
Erysimum_cheiranthoides_EcCYP87A126      IAPLICRKALKDIQYKNYTIPANWPVMVVPPAIHLDPNNYEDPLVFNPSRWEGS-EFTNA     407
                                         *.* * **.: : :        .* :**   ::*::   *::** *:* **:*  .   .

Calotropis_gigantea_cal_g015519.t1       SKNFMAFGGGMRLCVGADFAKLQMAIFLHHLVTKFRWTITHGGDTVRKPGLLFPNGLHVQ     470
Calotropis_procera_CpCYP87A103           SKNFMAFGGGMRLCVGADFAKLQMAIFLHHLVTKFRWTITHGGDTVRKPGLLFPNGLHVQ     470
Asclepias_curassavica_AC04g009170.1      SKNFMAFGGGMRLCVGADFAKVQMAIFLHHLVTKYRWTITHGGDTVRKPGLLFPNGLHVE     444
Asclepias_syriaca_AS04g025310.1.1        SKNFMAFGGGMRLCVGADFAKLQMAIFLHHLVTKYRWTITHGGDTVRKPGLLFPNGLHVE     470
Calotropis_gigantea_cal_g001177.t1       SKNFMAFGGGMRLCVGADFAKLQMAIFLHYLVTKYRWEITEGGNTIRKPGLLFPNGLHVH     435
Asclepias_curassavica_AC04g010230.1      SKNFMAFGGGMRLCVGADFAKLQMAIFLHYLVTKYRWVINGGGNTVRKPGLLFPNGLHVH     484
Asclepias_curassavica_AC04g010240.1      SKNFMAFGGGMRLCVGADFAKLQMAIFLHYLVTKYRWVINGGGNTVRKPGLLFPNGLHVH     484
Asclepias_syriaca_AS04g027770.1.1        SKNFMAFGGGMRLCVGADFAKLQMAIFLHYLVTKYRWVINEGGNTVRKPGLLFPNGLHVH     478
Asclepias_syriaca_AS04g027730.1.1        SKNFMAFGGGMRLCVGADFAKLQMAIFLHYLVTKYRWVINEGGNTVRKPGLLFPNGLHVH     478
Asclepias_syriaca_AS04g027600.1.1        SKNFMAFGGGMRLCVGADFAKLQMAIFLHYLVTKYRWVINEGGNTVRKPGLLFPNGLHVH     471
Asclepias_syriaca_AS04g027640.1.1        SKNFMAFGGGMRLCVGADFAKLQMAIFLHYLVTKYRWVINEGGNTVRKPGLLFPNGLHVH     476
Oryza_sativa_OsCYP87A6                   TKHFMAFGGGLRFCVGTDLSKVLMATFIHSLVTKYSWRTVKGGNIVRTPGLSFPDGFHIQ     509
Digitalis_purpurea_DpCYP87A106           SKTFMAFGGGMRLCIGADFTKVQMAVFLHCLVTKYKWKTIKGGDIVRCPGLKFPNGFHVQ     469
Digitalis_lanata_DlCYP87A4               SKTFMAFGGGMRLCIGADFTKVQMAVFLHCLVTKYKWKSIKGGDIVRCPGLKFPNGFHVN     472
Calotropis_gigantea_cal_g005561.t1       TRNFMAFGGGMRFCVGTDFTKVQMAVFLHCLVTKYKQGILMARPVLRADILQAARI----     464
Asclepias_syriaca_AS06g010210.1.1        TRNFMAFGGGMRFCVGTDFTKVQMAVFLHCLVTKYKWQETKGGDILRTPGLQFPNGFHVK     486
Asclepias_curassavica_AC06g015900.1      TRNFMAFGGGMRFCVGTDFTKVQMAVFLHCLVTKYKWQETNGGDILRTPGLQFPNGFHVK     489
Solanum_lycopersicum_SlCYP87A21          TRNFMAFGGGMRFCVGTDFTKVQMAVFLHSLVTKYRWQTIGGGNTVRTPGLQFPNGYHVR     467
Nicotiana_benthamiana_NbCYP87A122        SRNFMAFGGGMRFCVGTDFTKVQMAVFLHSLVTKYRWETIQGGDTLRTPGLQFPNGYHIR     467
Arabidopsis_thaliana_AtCYP87A2           SKHFMAFGGGMRFCVGTDFTKLQMAAFLHSLVTKYRWEEIKGGNITRTPGLQFPNGYHVK     466
Erysimum_cheiranthoides_EcCYP87A126      SKKFMAFGGGMRFCIGTDFSKLQTAVFLHSLLTKYSWEHISGGNMLRSPGLQFPNGYHVK     467
                                         :: *******:*:*:*:*::*:  * *:* *:**:      .    *   *
```

```
Calotropis_gigantea_cal_g015519.t1       ISAIKAN-----     477
Calotropis_procera_CpCYP87A103           ISAIKAN-----     477
Asclepias_curassavica_AC04g009170.1      ISAIKAN-----     451
Asclepias_syriaca_AS04g025310.1.1        ISTIKAN-----     477
Calotropis_gigantea_cal_g001177.t1       VINKPNAA----     443
Asclepias_curassavica_AC04g010230.1      VTEKPTAAIA--     494
Asclepias_curassavica_AC04g010240.1      VTEKPTAAIA--     494
Asclepias_syriaca_AS04g027770.1.1        VTDKPTAA----     486
Asclepias_syriaca_AS04g027730.1.1        VTDKPTAA----     486
Asclepias_syriaca_AS04g027600.1.1        VTDKPTAA----     479
Asclepias_syriaca_AS04g027640.1.1        VTDKPTAA----     484
Oryza_sativa_OsCYP87A6                   LFPKN-------     514
Digitalis_purpurea_DpCYP87A106           MSEREANQKACK     481
Digitalis_lanata_DlCYP87A4               MTERG-------     477
Calotropis_gigantea_cal_g005561.t1       ------------     464
Asclepias_syriaca_AS06g010210.1.1        ISEKVEDAREST     498
Asclepias_curassavica_AC06g015900.1      ISEKVEDAREST     501
Solanum_lycopersicum_SlCYP87A21          ISEKDEKIL---     476
Nicotiana_benthamiana_NbCYP87A122        LSEKDEKIQ---     476
Arabidopsis_thaliana_AtCYP87A2           LHKKRD------     472
Erysimum_cheiranthoides_EcCYP87A126      INKKEI------     473
```

**Figure S5: Alignment of CYP87A proteins.** The protein sequence alignment was generated by Clustal Omega. A phylogenetic tree of these sequences is shown in Figure 9B.
